# Standardizing carabid pitfall catches for trapping effort; a meta-analysis

**DOI:** 10.1101/2020.07.23.216994

**Authors:** Pavel Saska, David Makowski, David Bohan, Wopke van der Werf

**Affiliations:** Functional Diversity Group, Crop Research Institute, Drnovská 507, Prague 6 – Ruzyně, 161 06, Czech Republic; INRAE, UMR 518 AgroParisTech, INRAE, Université Paris-Saclay, 16 rue Claude Bernard F-75231 Paris Cedex 05, France; Agroécologie, AgroSup Dijon, INRAE, Univ. Bourgogne, Univ. Bourgogne Franche-Comté, F-21000 Dijon, France; Centre for Crop Systems Analysis, Wageningen University and Research, P.O. Box 430, 6700 AK Wageningen, The Netherlands

**Keywords:** activity-density, Carabidae, catch per unit effort, decline in diversity, pitfall traps, sampling effort, trap days, trappability

## Abstract

1. Carabid beetles (Coleoptera: Carabidae) provide important ecological services and are frequently used as a bio-indicator in monitoring environmental quality. The abundance and diversity of carabids is usually determined using pitfall trapping, but trap catches are difficult to compare between studies due to variation in trapping effort. The standardization of the catch for trapping effort has not been previously addressed in a global analysis of studies in the literature.
2. The aims of this study are (i) to define a method for estimating the effect of trapping effort on the size of the pitfall catch, and (ii) to explore factors related to study designs, sampling method, study origin, and level of data aggregation to determine how these factors affect the catch per unit effort in pitfall trapping.
3. We conducted a meta-analysis on the activity-density and diversity of carabids across studies, based on published data from Europe and North-America to analyse whether standardization of catch measurements might be possible. Data were extracted from 104 publications, spanning a period of 42 years.
4. The total catch was proportional to the number of trap days, and ranged from 0.19-9.53 beetles/(trap day) across studies (95% range), with a mean of 1.33 beetles/(trap day). The number of species was allometrically related to the trapping effort defined as the product of the number of traps, their perimeter and the time of exposure in the field, and characterized by a power exponent of 0.25. Species richness ranged across studies from 2.30-13.18 species/(m day)^0.25^ (95% range) with a mean of 7.15 species/(m day)^0.25^. The size of the catch and the number of species were higher in crops with narrow as compared to wide rows. There was no significant change in abundance or diversity of carabids in arable land over the 42 years covered. We also found that increasing trapping effort may not yield more accurate results.
5. The results show that it is possible to standardize activity-density-based catches and species diversity for trapping effort across studies using a power transformation, allowing meta-analysis of such data, e.g. to elucidate factors affecting abundance and diversity of the focal taxa.

## Introduction

Pitfall traps are commonly used to sample ground-dwelling invertebrates such as beetles or spiders. The traps are typically plastic or glass containers sunk into the soil. Pitfall catches cannot be directly interpreted as abundance estimates because the number of trapped individuals depends not only on their population density but also on their activity (particularly movement speed) and behaviour, e.g. a preference for movement on vegetation or soil, and behaviour near the edge of the traps (Adis 1979). Movement speed is affected by extrinsic biotic factors, e.g. resistance of the vegetation (Heydemann 1957), as well as intrinsic biotic factors, such as the body size (Engel *et al.* 2017), level of hunger (Wallin & Ekbom 1994) or sexual motivation (Mols 1993), and abiotic factors, e.g. temperature (Saska *et al.* 2013). The “trappability” may also differ between species according to their behaviour, e.g. movement speed or whether individuals are able to avoid falling over the edge of the trap (Halsall & Wratten 1988) or to escape (Luff 1975). Thus, the pitfall catch is usually treated as a measure of “activity-density” (Adis 1979). There is a consensus in the field that the difficulty of relating the pitfall catch to the density and species richness of invertebrate stocks is an impediment to the interpretation of field studies based on pitfalls (Kotze *et al.* 2011).

Estimating true densities of populations based on sampling has been a continuing topic in ecology (Southwood & Henderson 2000; Skalski, Ryding & Millspaugh 2005). In fisheries, the true abundance of a fish stock can be assessed from catch data assuming a relationship between the catch (*C*), the abundance of the individuals (*N*), the fishing effort (*E*) and a factor “catchability” *q* that is related e.g. to animal behaviour and its spatial distribution (Harley, Myers & Dunn 2001; Martell 2008):

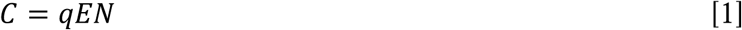

with underlying error structure to be log-normally distributed; the catch per unit effort, 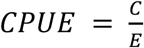, is assumed to be proportional to population size (Harley, Myers & Dunn 2001; Martell 2008). In this paper we extend this approach to invertebrate pitfall trap catches, conducting a meta-analysis of published data on the pitfall-trap-based catches of carabid beetles (Coleoptera: Carabidae).

We chose carabid beetles as a model group of organisms because they have been recognized as important biocontrol agents in agricultural crops, preying upon invertebrate pest species (Sunderland 2002) and eating and potentially regulating seeds of arable weeds (Bohan, Boursault, Brooks & Petit 2011; Frei, Guenay, Bohan, Traugott & Wallinger 2019). Because of their important contribution to these ecological services, conservation and augmentation of stocks of carabids in farmland is a primary concern of agro-ecological management (Brooks *et al.* 2012). Carabids also represent an excellent model for environmental monitoring and bio-indication because they respond to abiotic and biotic variation, and to disturbances and management (Kotze *et al.* 2011).

This paper addresses two aims. First, we aim to define a method for estimating the effect of trapping effort on the size of the pitfall catch in order to allow a rigorous analysis of the effects of environmental and management factors on the catch. Second, we explore factors related to study designs, sampling method, study origin, and level of data aggregation to determine how these factors affect the catch per unit effort in pitfall trapping. Kotze *et al.* (2012) demonstrated that the effort of trapping, particularly where traps are lost or damaged, should be taken into account in evaluating pitfall catches. The question of how best to generalise and integrate trapping effort in a meta-analysis of studies with many different trap designs and layouts, remains to be addressed.

Finding a way to control the effect of trapping effort on the catch would make comparison of pitfall catches across studies possible, thus rendering useful the enormous corpus of published information that is currently inaccessible to systematic comparison, e.g. by meta-analysis. We are aware of no previous papers on the topic of standardizing pitfall trap catches for trapping effort across studies. We therefore conduct a meta-analysis of published literature and explore possible proxies for trapping effort that have a good relationship to the size and diversity of the catch.

## Materials and methods

### Conceptualising the catch per unit effort of pitfall trapping

*A priori*, we expect an allometric relationship between the pitfall catch size, *C*, and trapping effort, *E*:

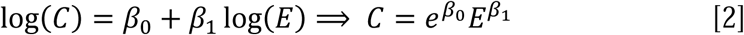

where *C* can represent the total number of individuals caught (total catch). If *β*_1_ = 1, the relationship between the catch and trapping effort is linear and the constant 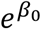 would represent the product of catchability (or better trappability if pitfall traps are considered) and abundance (*qN*) of equation [1]. An analogous equation can be constructed for the species richness in the catch, *S*:

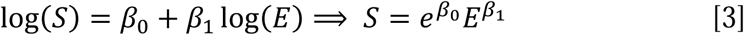

The constant 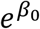 from eq. 2 and 3 is the number of individuals or species caught per unit of effort *E*, raised to a power *β*_1_.

We consider several possible proxy variables to characterize trapping effort *E*: number of traps (*K*) (Kotze et al. 2012), total trap perimeter (*P*) (Turin *et al.* 1991), duration of trap exposure (*X*) (Jung, Jeong & Lee 2019), trap-days (*R*), i.e. the product of *K* and *X* (Kromp 1989), and a new metric - perimeter-days (*Q*), i.e. the product of *P* and *X*.

### Search strategy and data

Searches were made in the Science Citation Index Expanded (SCI-E) and Scopus for the years 1945-2018 (Appendix S1). Search #1 was made in SCI-E and covered the years of 1991-2018, and used the following search string: (carabid* OR “ground beetle*”) AND (field* OR crop*) AND (*icide* OR manag* OR control* OR organic* OR conventional* OR practice* OR cultivation OR till*) AND (“species richness” OR “number of species” OR diversity OR activity* OR abundan*) NOT (wood* OR forest* OR vineyard* OR olive* OR orchard* OR urban* OR wetland* OR highway*). Search #2 was also made in SCI-E. It covered the years 1945-1990 and was less restrictive, because abstracts were not available for articles before 1991: (carabid* OR “ground beetle*”) AND (*icide* OR manag* OR control* OR organic* OR conventional* OR practice* OR cultivation OR till*) NOT (wood* OR forest* OR vineyard* OR olive* OR orchard* OR urban* OR wetland* OR highway*). Search #3 was made using Scopus (Elsevier) and covered the years 1960-2018: (carabid* OR “ground beetle*”) AND (field* OR crop*) AND (*icide* OR manag* OR control* OR organic* OR conventional* OR practice* OR cultivation OR till*) AND (“species richness” OR “number of species” OR diversity OR activity* OR abundan*) AND NOT (wood* OR forest* OR vineyard* OR olive* OR orchard* OR urban* OR wetland* OR highway*). After removal of duplicates, this search resulted in 648 publications. Then, titles, keywords, abstracts and full text were screened retaining only those papers containing primary data on pitfall trapping of carabid beetles and including information on field and crop management. The final database comprised data from 104 publications (Appendix S2) and 810 data records.

The resolution of data extraction followed the level of aggregation over experimental units and over seasons in the source publication. Studies based on replicate plots (“plots”) reported means or averages over the plots. Studies in which the single field was the experimental unit, the data were either aggregated over the multiple sampled fields (“multiple fields”), or were available for each single field (“single fields”). In each of those categories, there were studies that reported data for each season separately and studies that aggregated data over multiple seasons. Altogether, we considered six different types of studies when considering data aggregation over both units and seasons (Appendix S3).

Information on the number of carabid beetles caught was available from 101 publications. For these studies, we calculated n = 792 values of the total number of carabids caught (*C*). The total number of species (*S*) was extracted from 49 publications (n = 335). Variables related to study design and trapping effort were extracted from each publication (Table 1). The total perimeter of the traps (Turin *et al.* 1991) was calculated as *P =* π*dK* for circular traps where *d* is trap diameter (cm), and *K* is the number of traps, or as *P = 4lK* for square traps where *l* was the side length. The number of trap-days (*R*) was calculated as *R = XK* where *X* is the number of days of exposure, and perimeter-days (*Q*) were calculated as *Q = XP*.

**Table 1.**
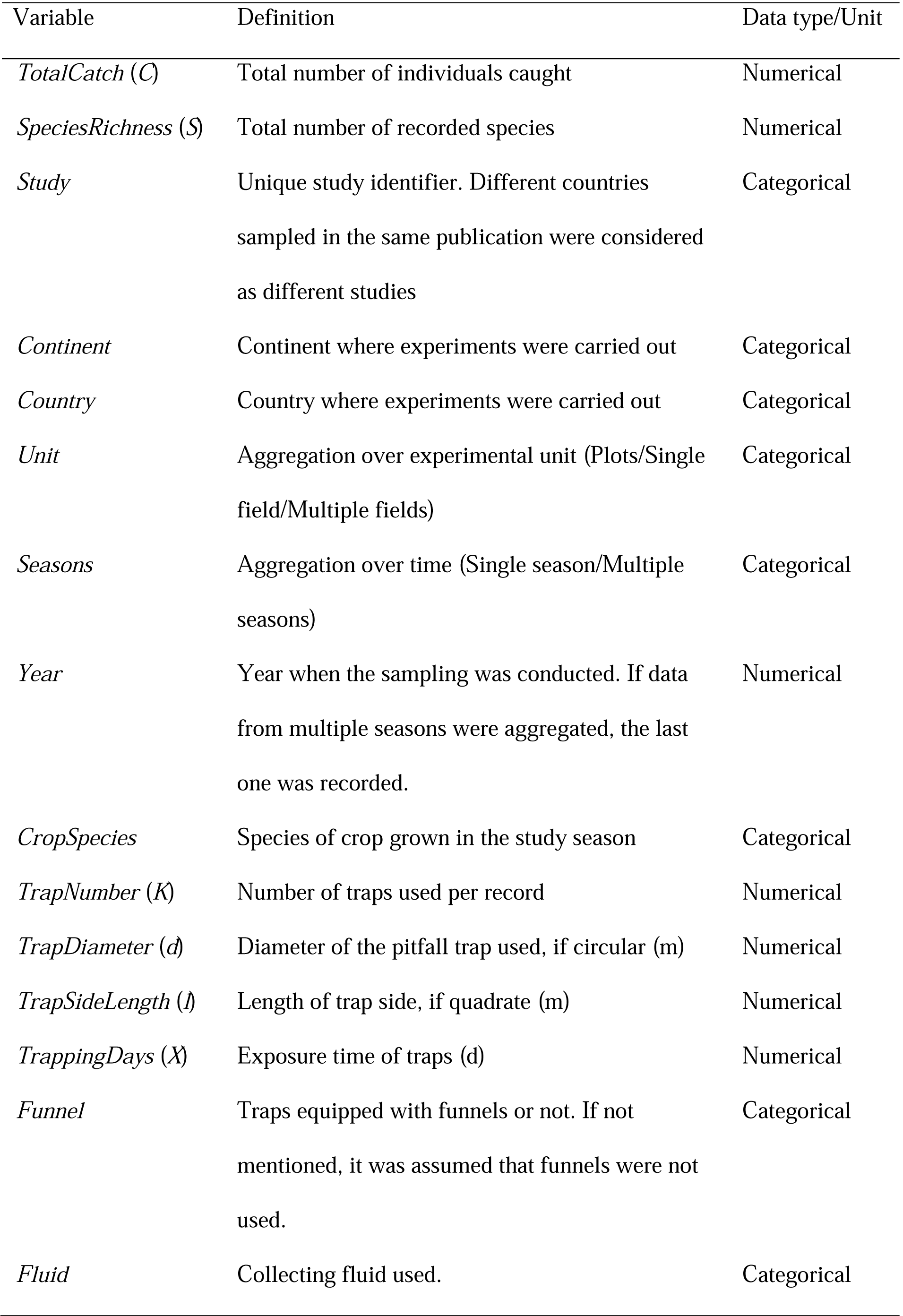
Data extracted from publications

### Statistical analysis

In a first step of the analysis, we compared five different variables measuring trapping effort (see above) to standardize the total catch (*C*) and species richness (*S*). Based on *a priori* testing that included Gaussian, Poisson and negative binomial distributions, we used the Gaussian error structure and identity link (having both dependent and independent variables on the log scale) for total catch and the Poisson error structure with log-link for species richness (Appendix S4, Table S1). We selected as the best proxy for trapping effort the measure that was the most closely related to the total catch or species richness, as assessed by information criteria (AIC, BIC) and by the precision of the slope estimate of the model relating total catch or species richness to trapping effort. The best model was used to estimate the value of *β*_1_ (used in Eqs. [2-3]). All analyses used mixed effects models in which random factors were included to account for effects of study and year of sampling within a study (Appendix S4, Table S1). Independent variables were centred to remove correlation between the slope and the intercept and to increase the robustness of fitting. Since there was no measure of the variance available for the total catch or species richness in the source studies, we used the log of the measure of trapping effort as weights (Appendix S4, Table S1). The adequacy of including weights in the preferred models was assessed with AIC and BIC. All analyses were performed in R 3.6.1 (R Core Team 2019), and mixed effects models were fitted using lmer (total catch) or glmer (species richness) functions of the package lme4 (Bates, Machler, Bolker & Walker 2015).

In the second step of the analysis, we explored the effects of the variables related to study design and data aggregation in the reporting of the results. We used as predictors the year of sampling (*Year*), continent (*Continent*), crops grouped in two categories according to row width (*RowWidth*), presence of a funnel (*Funnel*), type of collecting fluid (*Fluid*) and level of aggregation over experimental units (*Unit*) and over experimental seasons (*Season*) (Table 1). The effect of these variables was tested separately by lmer/glmer. In order to take the effect of *E* into account, we included 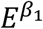 as an offset (Kotze et al. 2012) in the models relating the observed values of *C* and *S* to the considered factors (Appendix S4, Table S2). Because of missing data for *RowWidth* and *Fluid* in several records, we used a reduced data set in analyses including these variables (n = 721 for individuals and n = 305 for species). The effect of the categorical variables (*Unit, Season, Continent, RowWidth, Funnel, Fluid*) was further assessed by comparing the cumulative distributions of the effort-adjusted catch and species richness, 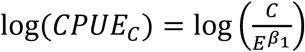 and 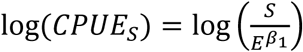, respectively, for the groups of records using the Kolmogorov–Smirnov test.

Given that the predictor variables of the study design and aggregation in reported results are likely to be correlated, we used the multi-model inference approach (Burnham & Anderson 2002) to assess the importance of these predictors. We formulated global models; one for the number of individuals caught and one for the observed species richness (Table S3 in Appendix S4), and used the function dredge (R package **`MuMIn`**; Barton 2009) to rank the models. Random effects, weights and error structure were derived from the best models describing the relation between the catch with unit effort, including an offset related to sampling effort (defined as explained above). Fitted models were automatically ranked according to AICc and BIC, and the set of top models was delineated by 6 units of AICc or BIC, respectively (Grueber, Nakagawa, Laws & Jamieson 2011). Model averaging revealed the relative importance of explanatory variables based on the top models, along with the relationship between response and explanatory variables (Burnham & Anderson 2002), and was performed using the function model.avg (R package **`MuMIn`**; Barton 2009). The parameters of the averaged model and their standard errors were estimated using the zero-method which calculates the weighted mean coefficient estimates over the selected models substituting a zero if a predictor was not selected in a model (Grueber, Nakagawa, Laws & Jamieson 2011). Marginal R^2^ values indicate the amount of variation explained by fixed factors only, while conditional R^2^ values represent the variance explained by both fixed and random factors (Nakagawa & Schielzeth 2013).

### Publication bias

Publication bias was assessed by constructing funnel plots for the log(*CPUE*_*C*_) and log(*CPUE*_*S*_). We used the log of the optimal expression of 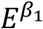 as the measure of study precision assuming that data records with exerted greater trapping effort will provide more precise estimate of the *CPUE*_*C*_ and *CPUE*_*S*_. There is no publication bias if the data points are symmetrically spread over the left and right side of the triangle. Points outside the funnel indicate possible outliers or heterogeneity in the data. Publication bias was assessed using all available data records.

## Results

### Descriptive analysis

Of the 104 studies, 68 originated from Europe and 36 from North America. Altogether the data came from 22 countries. Most publications came from the USA, Germany and Canada.

Most records are based on single sampling season (n = 726); much fewer records were based on multiple seasons (n = 84) (Figure 1a). There were slightly more records from Europe (n = 451) than from North America (n = 359). Most of the data records came from plot experiments (n = 604), and fewer records were based on single fields (n = 116) or multiple fields (n = 90). Plot-based studies were more frequent in North America (ca. 87 % of records) than in Europe (ca. 63 %) (Figure 1a).

**Figure 1.**
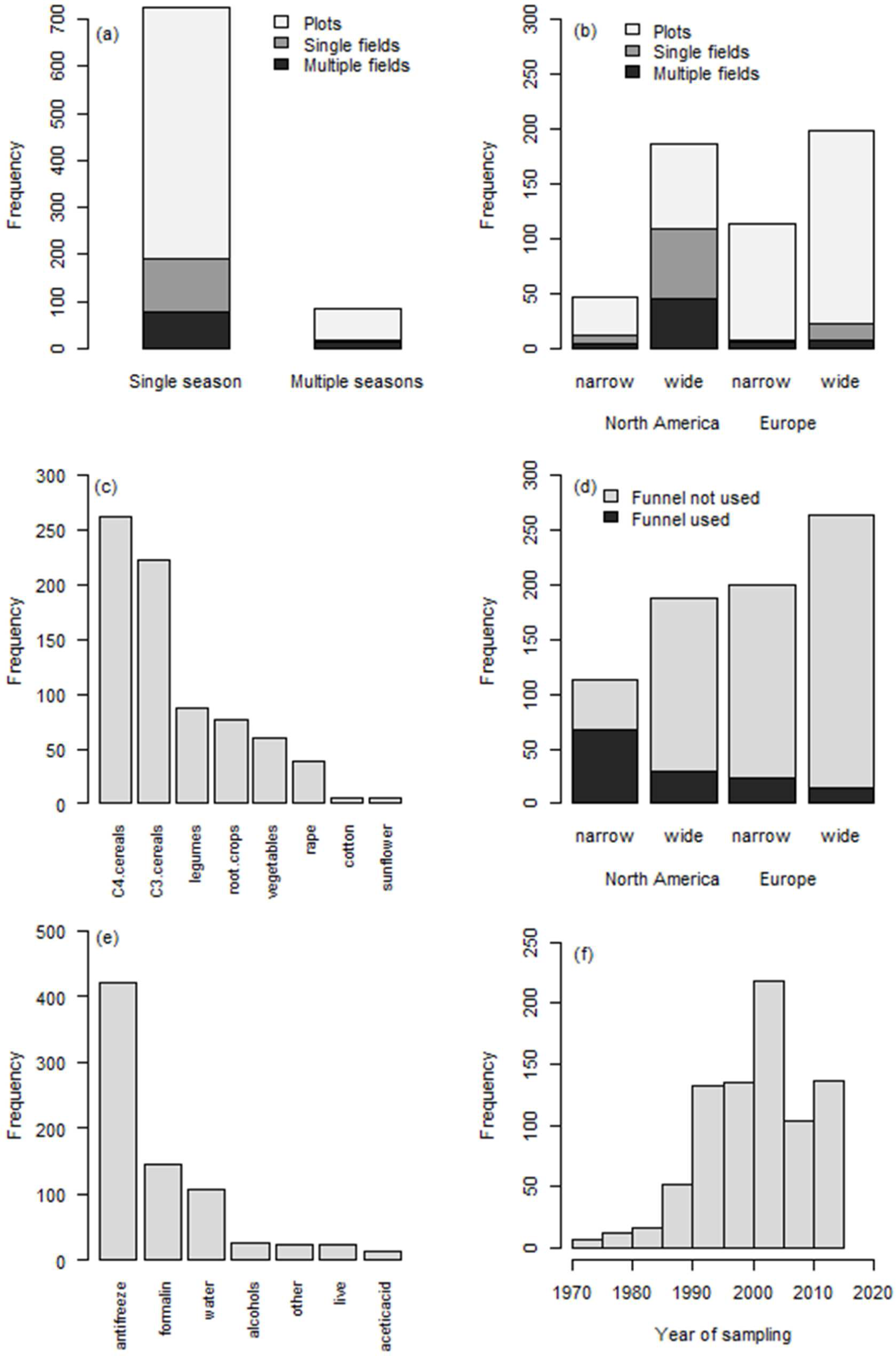
Distribution of records in the data set with variables fundamental to the study design, sampling scheme and level of data aggregation in the source publications. (a) Number of data records according to aggregation over experimental units and sampling seasons. (b) Number of data records originating from North America and Europe, from crops with wide or narrow rows, and from plots, single fields, or multiple fields. (c) Number of data records related to experimental crops. C3 cereals include wheat (n = 141), barley (66), oats (3), triticale (2) and undefined cereals (10); C4 cereals include both sweet corn and corn for silage (260) and sorghum (2); legumes include pea (35), soybean (24), beans (18), lupin (6) and faba bean (4); root crops include potatoes (41), sugarbeet (31) and carrots (5); and vegetables consist of tomatoes (16), squash (10), melon (8), cauliflower (8), cabbage (7), zucchini (6) and onion (5). Cotton (6), oilseed rape (10) and sunflower (5) are single crop categories. (d) Number of data records based on trapping with or without funnels, and originating from North America or Europe, and from crops with wide or narrow rows. (e) Number of data records related to the collecting fluid used. Antifreeze include ethylene glycol (236), propylene glycol (171) and unspecified antifreeze (14), water includes water with (66) or without (41) diluted salt, alcohols include ethanol (17) and iso-propyl alcohol (8), other include Na_3_PO_4_ (4), CuSO_4_ (12), natrium benzoate (2) and unspecified liquid (6), and acetic acid include mixtures that contain this ingredient (14). Formalin (144) and live traps without any collecting fluid (23) are single type categories; (f) Distribution of data records according to the year of sampling.

The experimental crops were unevenly represented in the data set (Figure 1c), with C4 and C3 cereals dominating. Records for wide row crops were more frequent (n = 463) than for narrow row crops (n = 300). Crops with wide row spacing are more common in North American than in European studies (Figure 1a). In Europe, wide row crops were more frequent in plots than in single of multiple fields (Figure 1b).

Studies which used funnels in the traps were less frequent (n = 140) than those without (n = 670). Funnels were more frequently used in North America (ca. 26 % of records) than in Europe (10 % of records) (Figure 1d). Altogether 16 different collecting fluids were used in this data set which were grouped in six categories (Figure 1e). Traps to collect live beetles formed an additional category (Figure 1e). The data set is dominated by fluids based on antifreeze compounds, followed by formaldehyde and water (usually containing diluted salt) (Figure 1e). Funnels were used only in traps that contained antifreeze compounds, alcohols and a CuSO_4_ solution.

Data records originated from a period of sampling spanning 43 years, from 1973 to 2015, but most records (62 %) came from studies conducted between the years 2000 and 2015 (Figure 1f).

Variability in continuous input variables related to trapping effort, total catch and observed species richness is shown in Appendix S5.

### Finding the optimal standardization of total catch and species richness per unit effort

The total catch significantly increased with all measures of trapping effort. Trap-days, *R*, was superior to all other alternative measures of trapping effort (Appendix S6). Each of the criteria used for model comparison identified different models as best. We chose model C18 as our most preferred model (Fig. 2a) because it estimated the slope parameter *β*_1_ with greater accuracy than model C20 (Appendix S6). The final model for standardizing the catch per unit effort (C27) differed from the model C18 by centring the trap-days *R* and using weights based on the trap-days (ΔAIC = 16.8). The estimated slope value of this best model C27 was *β*_1_ = 0.959 ± 0.056 which was not significantly different from 1 (*P* = 0.471), indicating that the number of individuals caught is proportional to the trapping effort expressed as trap-days, and the effort-adjusted catch is equivalent to 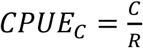. The mean *CPUE*_*C*_ across the entire data set was 1.33 ± 0.12 individuals per trap day.

**Figure 2.**
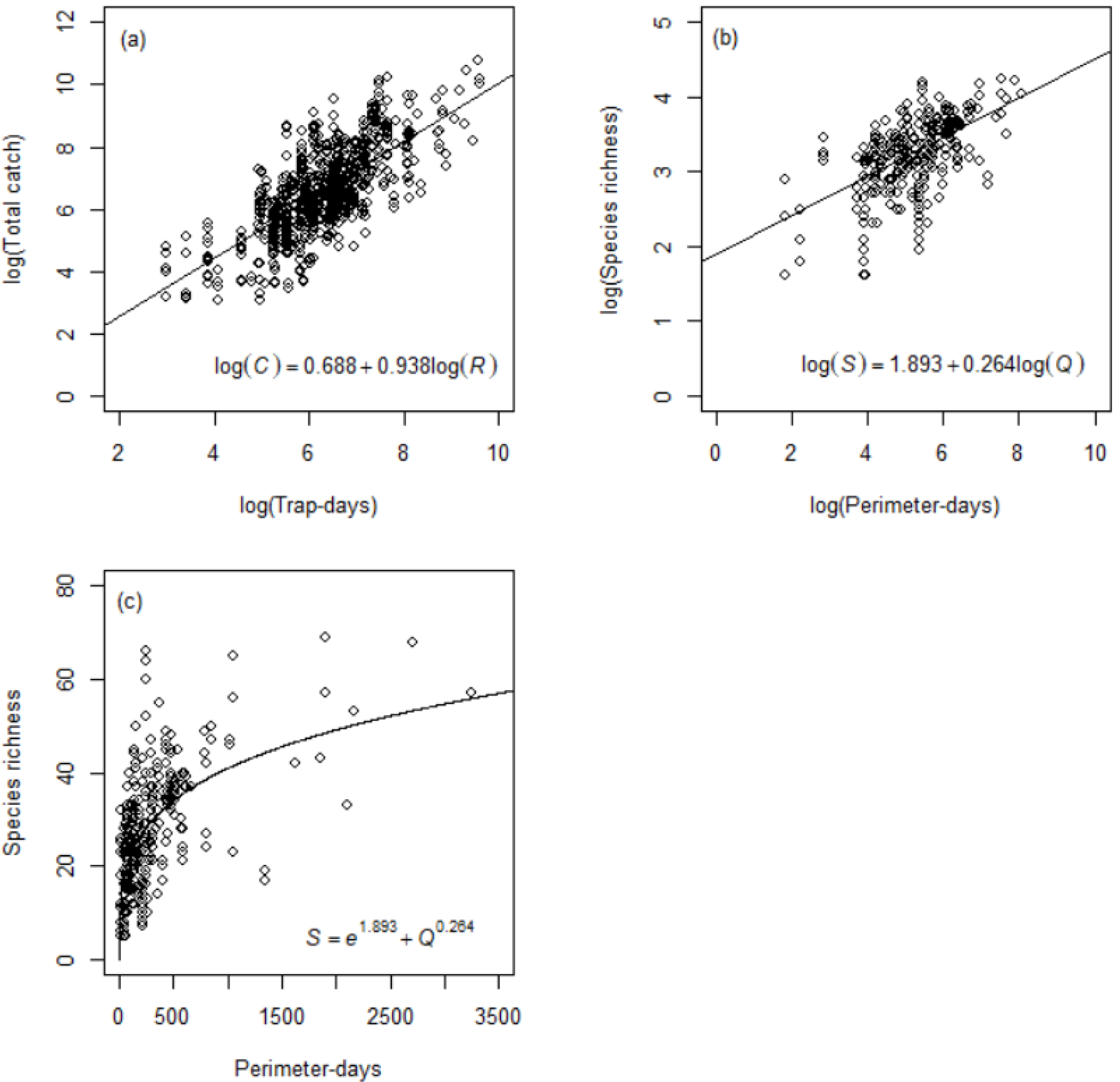
Dependency of carabid pitfall catch on trapping effort. (a) The effect of trap-days (*R*) [log(trap days)] on the total catch [log(individuals)] (model C18; Appendix S6); (b) The effect of perimeter-days (*Q*) [log(m days)] on the observed species richness *S* [log(species)] (Model S23; Appendix S6); (c) same as (b) but on the natural scale.

Species richness increased significantly with all measures of trapping effort, and perimeter-days *Q* was the most effective measure of standardization (Appendix S6). A random intercept model, S23, was identified as best (Appendix S6), and was further improved by centring the perimeter-days *Q* (S26); adding weights according to the perimeter-days was not justified (ΔAIC = 573.8). The estimated value of the slope parameter in model S26 was *β*_1_ = 0.257 ± 0.037, indicating that the number of recorded species increases substantially less than proportionally with *Q* (Fig. 2b-c). The effort-adjusted species richness is therefore equivalent approximately to 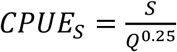. Mean *CPUE*_*S*_ across the entire data set was 7.15 ± 0.37 species (m d)^-0.25^.

### Factors influencing effort-adjusted catch size and species richness

Effort-adjusted catch significantly varied with *Continent, RowWidth* and *Fluid*, but not with *Unit, Season* and *Funnel* (Figure 3). The *CPUE*_*C*_ was on average 64.9 % higher in Europe than in the North America, and lower by 31.7 % in crops with wide rows compared to crops with narrow rows. Live traps provided 84.2% reduction in *CPUE*_*C*_ compared to traps containing fluids based on acetic acid that assured the highest *CPUE*_*C*_, or 43.0% reduction compared to the most commonly used antifreeze-based fluid. The effort-adjusted species richness was affected significantly by the *RowWidth, Season* and *Funnel*, but not by *Continent, Unit* and *Fluid* (Figure 4). The *CPUE*_*S*_ was reduced by 28.9 % in crops with wide rows compared to narrow rows, by 23.0 % if data were aggregated over multiple seasons, but increased by 89.5 % if funnels were used inside the traps. The comparison of the cumulative probability distributions of *CPUE*_*C*_ and *CPUE*_*S*_ for categorical variables provided similar results (Appendix S7). Neither effort-adjusted catch nor species richness showed a significant temporal trend over the period covered by this study (Figure 5).

**Figure 3.**
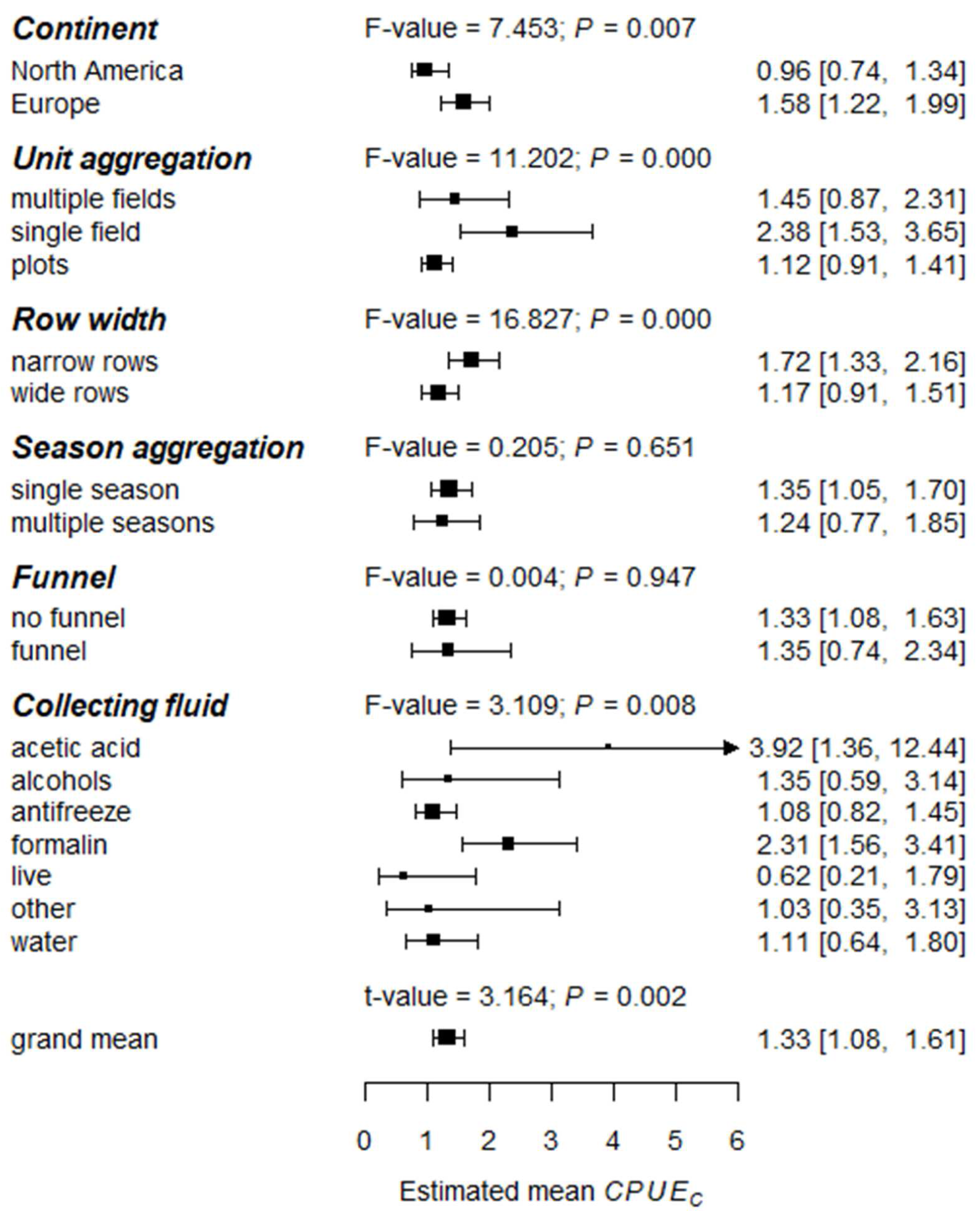
Estimated mean effort-adjusted catch, *CPUE*_*C*_ [individuals (trap day)^-1^], shown for particular categories of variables potentially affecting the relationship with the pitfall catch and trapping effort. Size of the symbols are relative to sample size. Horizontal bars are 95% CI bootstrapped by the bootMer function (R package lme4, 200 simulations).

**Figure 4.**
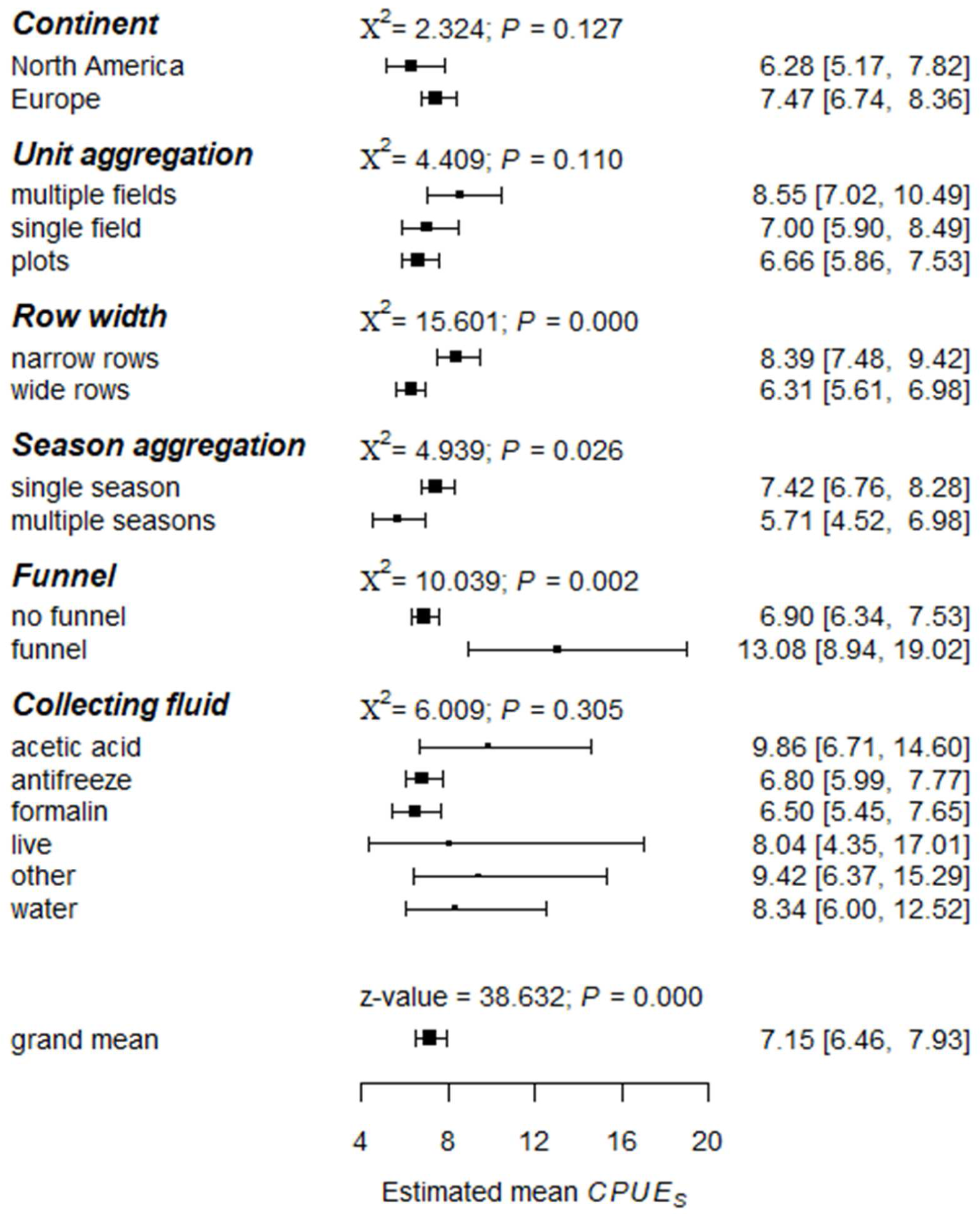
Estimated mean effort-adjusted species richness, *CPUE*_*S*_ [species (m day)^-0.25^], shown for particular categories of variables potentially affecting the relationship with the pitfall catch and trapping effort. Size of the symbols are relative to sample size. Horizontal bars are 95% CI bootstrapped by the bootMer function (R package lme4, 200 simulations).

**Figure 5.**
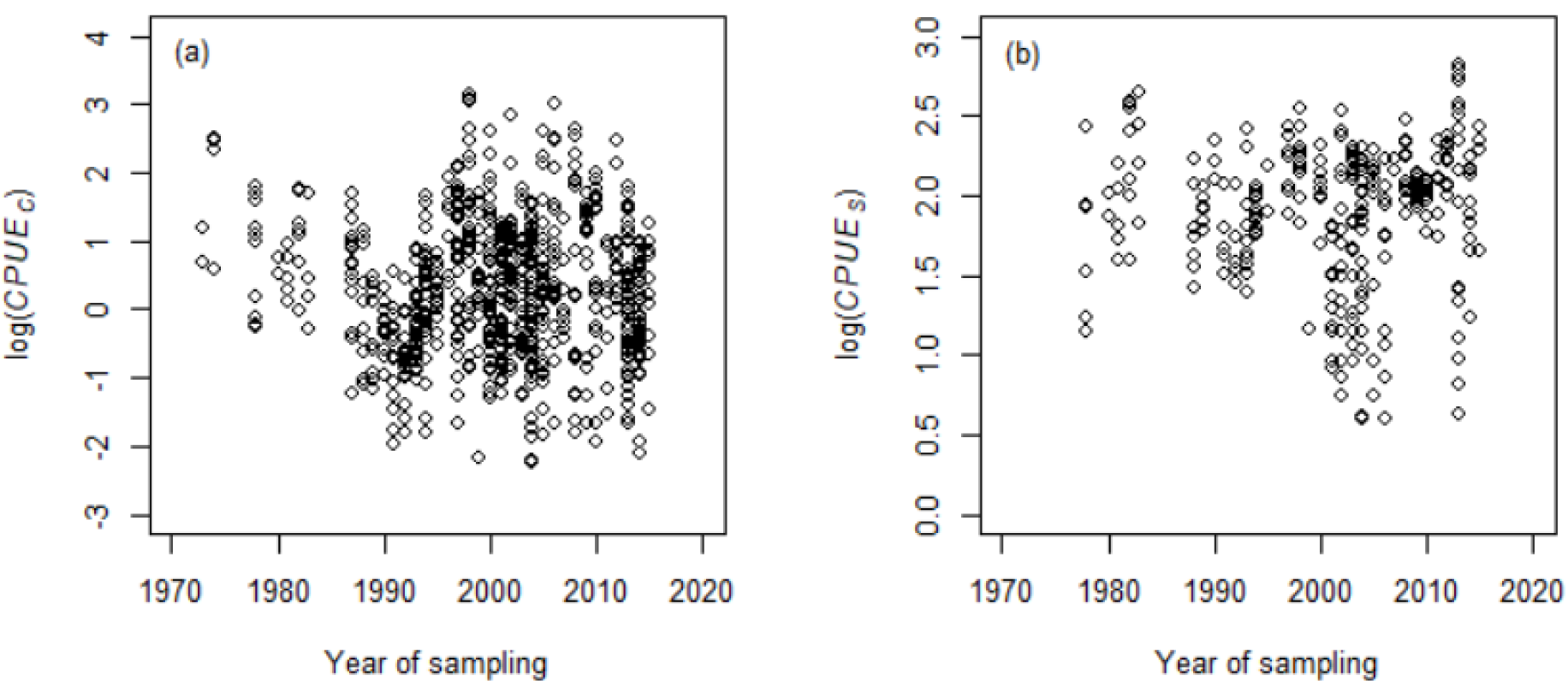
Variation in the effort-adjusted catch, *CPUE*_*C*_ [individuals (trap day)^-1^] (a), and effort-adjusted species richness, *CPUE*_*S*_ [species (m day)^-0.25^] (b) over the years of sampling covered by the data set.

The results of multi-model inference were different from the results of regression models using one variable at a time. Averaging the top models (Appendix S8) revealed that *RowWidth* was the only variable that significantly affected effort-adjusted catch, regardless of the information criterion used for model selection. The effect of other variables proposed by the single regression models (Figure 3) cancelled each other out, probably due to correlations between inputs, or the reduced data set used for multi-model inference. Using AICc, *CPUE*_*C*_ was by 30.4 % lower in crops with wide row span compared to narrow row span (z-value = 3.788, *P* < 0.001), which is close to the value estimated by the single variable regression model. The values estimated based on BIC weights were very similar to those calculated with AICc weights.

Effort-adjusted species richness was significantly affected by *RowWidth, Funnel* and *Season* if the model selection was based on AICc, and only by the *RowWidth* if the model selection was based on BIC. Crops with wide rows reduced the *CPUE*_*S*_ by 22.2 % (AICc; z-value = 3.564, *P* < 0.001; BIC-based selection gave very similar values), the use of a funnel increase the *CPUE*_*S*_ by 66.0 % (AICc; z-value = 2.603, *P* = 0.009), and aggregation of data over multiple seasons reduced the *CPUE*_*S*_ by 26.3 % (AICc; z-value = 2.075, *P* = 0.038). These values are also close to the results of single variable regression models (see above).

### Publication bias

Studies with trapping effort of less than 20 trap days (*R* = *e*^3^), or ca. 5.5 m d (*Q* = *e*^1.7^) were absent from the literature (Figure 6). The variability in the log(*CPUE*_*C*_) or log(*CPUE*_*S*_) did not change with trapping effort (Figure 6), which suggests that the variability is unrelated to the study precision and represents biological variation.

**Figure 6.**
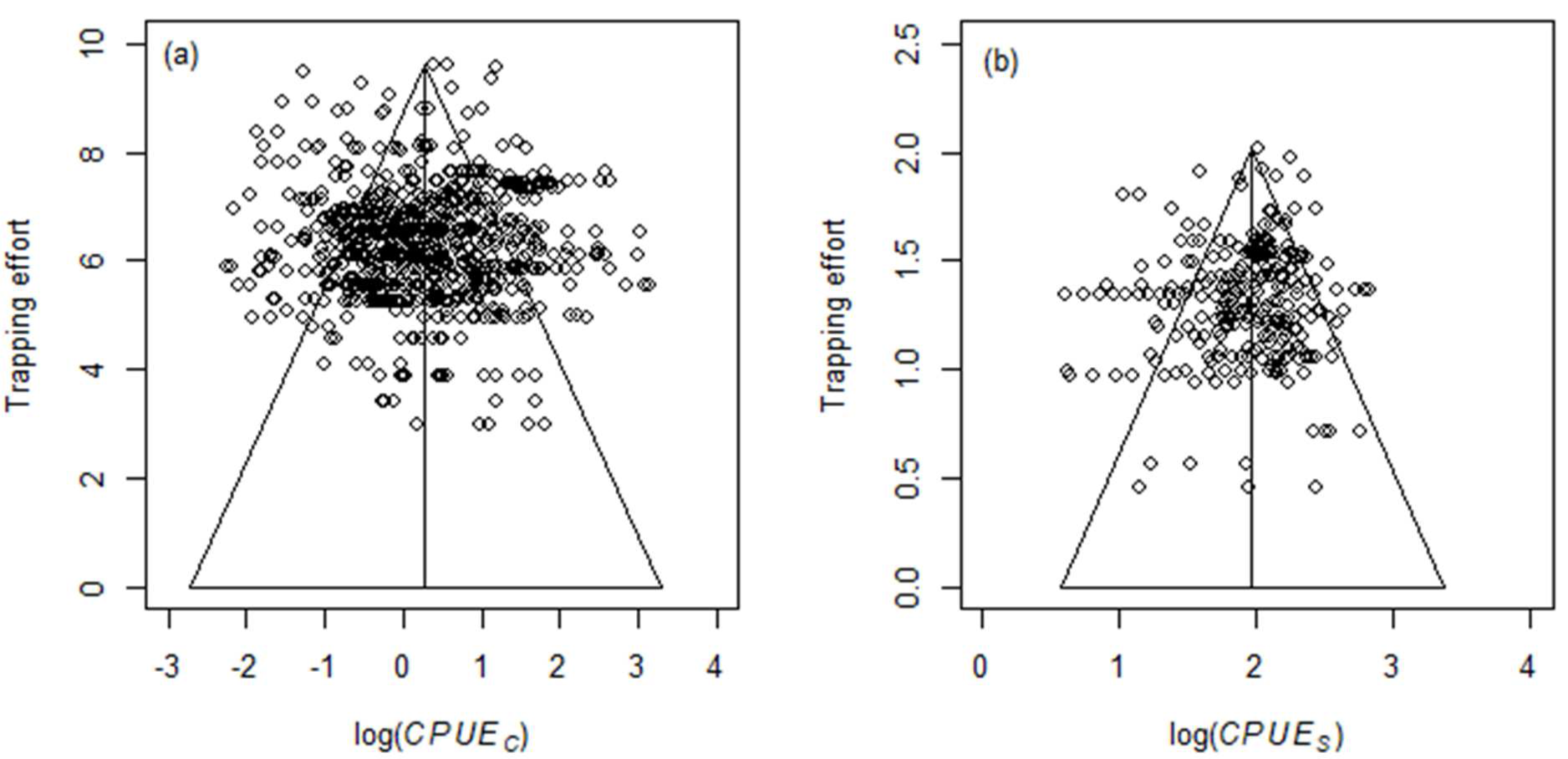
Assessment of publication bias in the data included in the meta-analysis, using funnel plots of the effort-adjusted catch [log(individuals (trap day)^-1^)] against trapping effort as trap-days [log(trap day)] (a), and the effort-adjusted species richness [log(species (m day)^-0.25^)] against trapping effort as perimeter-days [log((m day)^0.25^)] (b). The vertical lines represent the overall means on the log scale, as predicted by the null models.

## Discussion

This analysis showed that trap days was the most suitable measure for expressing trapping effort when analysing the number of carabid beetles caught in pitfall traps. The exponent of the allometric relationship was not significantly different from 1, indicating that the catch is proportional to trapping effort, without significant curvature or saturation in the relationship. The best measure to express trapping effort when analysing species diversity of the catch was the number of meter days to the power of 0.25, where the meters refer to total perimeter length of all traps while days refers to the time of trap exposure in the field. The exponent value of ca. 0.25 was significantly less than 1, indicating that the relationship between trapping effort and species diversity is strongly non-linear, with species diversity increasing quite slowly with increases in the number of perimeter of traps or the time of exposure.

We also found that increasing trapping effort may not yield more accurate results. Large heterogeneity across the records demonstrates the importance of the local conditions for determination of the actual effort-corrected catch and richness, and this is largely captured by the random terms of the models. This finding would also suggest that to obtain more information about arthropod communities in arable land, it is of greater importance to sample more locations and more different conditions, than to expend great trapping effort in any particular location or condition.

Although the single variable regression models identified multiple variables that affected the effort-adjusted catch and species richness, model averaging indicated that the contrast between wide and narrow row crops was the major factor in the overarching analysis. This discrepancy between the two approaches may be due to the fact that the influence of particular factors, though locally important, were confounded and thus cancel each other out in a global analysis. As a consequence, our study suggest that no concern is needed in a future (meta-) analysis about the variability in experimental design, sampling scheme or trap design across source studies.

Crops with wide or narrow rows differ in phenology and structure. Crops with wider rows (e.g. corn) constitute a less favourable environment, because there is greater proportion of bare soil that needs longer time to reach a closed canopy. Bare soil is unfavourable for many carabid species since the risk of predation increases (Eyre, Luff & Leifert 2013; Seidl, Gonzalez, Kadlec, Saska & Knapp 2020), and diurnal changes in microclimate are much more prominent than under a more closed crop canopy (Rosenberg, Blad & Verma 1983; Krédl, Středa, Pokorny, Kmoch & Brotan 2013). Crops with wider rows also allow crop management, such as mechanical weed control, which may be more intensive and extend longer over the season, potentially disturbing development stages of carabids present in soil.

The effort-adjusted species richness was also affected by the level of data aggregation over seasons (i.e. single season vs. multiple seasons) and the use of funnels inside the traps. The former can readily be explained by the fact that even though the local populations of carabids species show asynchronous fluctuations between seasons (Kotze *et al.* 2011), which results in a change in the relative contribution of particular species to the catch between years (e.g. Vesely & Sarapatka 2008), the number of newly recorded species per year on sites sampled for multiple seasons is low. Increasing the trapping effort over more seasons on the same site brings disproportionately fewer new species recorded than adding a new site, sampled with the same effort. The presence of funnels increased observed species richness, which may be associated with a reduction in the probability of escaping from the traps (Obrist & Duelli 1996). Interestingly, the use of funnels did not affect the number of individuals caught.

Remarkably, we were unable to detect trends in *CPUE*_*C*_ or *CPUE*_*S*_ over the 43 years covered by the data set, suggesting that carabid populations have not declined in abundance or diversity in arable fields on broader geographical scale over this period. This result is in contrast to the monitoring programmes on local (Poszgai, Baird, Littlewood, Pakeman & Young 2016; van Noordwijk *et al.* 2017) or national scales (Brooks *et al.* 2012; Ewald *et al.* 2015), which have found carabid populations to have declined over time, as well as with general perception that insect populations decline in terrestrial ecosystems (Eggleton 2020; van Klink *et al.* 2020). Since the present data set originates from many local independent studies performed in different years, it is less sensitive to site-specific inter-annual fluctuations (Kotze *et al.* 2011). Biological explanation for the lack of any trend might be that species inhabiting annual arable fields are adapted to “early successional stages” of vegetation development that arable fields are in fact representing, and to periodical disturbances due to management measures within the fields. Given that local declines of carabids, as well as other insects, have been observed in non-crop semi-natural habitats (Brooks *et al.* 2012; Ewald *et al.* 2015; Hallmann *et al.* 2017) a reasonable expectation would be that this decline should spill over into arable fields as many carabid species recolonize fields from these non-crop habitats (Tscharntke, Rand & Bianchi 2005). The present data, however, do not provide support for this assumption. Evaluating if the community composition of the study group has changed in arable fields over the years was not, however, possible on our data because this evaluation would require different data extraction procedures than we have employed.

Our results do beg the question: “Can the newly established relationships for *CPUE* help us to relate the size of the pitfall catch to the real densities or diversity of carabid population in the field?” Not on its own, but since we resolved one of variables from the relationship between the catch size and density (eq. 1; Harley, Myers & Dunn 2001; Martell 2008), i.e. trapping effort, we have advanced closer to a reliable approximation of the field densities and diversity from the pitfall trap catches. What remains to be investigated is quantification of trappability coefficients *q* for a range of species, because the likelihood of being trapped is likely to be species specific (Halsall & Wratten 1988; Engel *et al.* 2017). Thus, eq. 2 can be further extended to consider species specific trappability coefficient *q*_*i*_. A relationship for *CPUE*_*C*_, with species-specific catches *C*_*i*_ and trapping effort expressed as trap-days *R* would be:

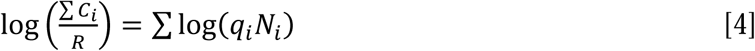

Obtaining the trappability coefficients *q*_*i*_ for a range of species will be needed, e.g. within enclosures or for manipulated densities, from various environments and in different abiotic conditions. The framework presented here may serve as a tool to get closer to the reliable estimates of population densities, using pitfall traps.

In this paper we establish relationships for the catch of carabid beetles, expressed as the number of individuals and the number of species, as a function of the unit effort of pitfall trapping. This method that we have applied to carabid beetles, could similarly be applied to other taxa with similar ecology, trapped using pitfalls, including ectothermic vertebrates or small mammals, or for other trapping devices that give activity-density estimates, such as window or suction traps used for collecting flying insects. Standardization of the pitfall catch for trapping effort will be very useful in future systematic comparisons of multiple independent catches, since the data collected in various conditions are thus made more comparable. This approach removes one of the obstacles that has hampered meta-analyses of pitfall trap data.

## Acknowledgements

This work was conducted within an ERA-NET C-IPM project BioAWARE. The stay of P.S. at Wageningen University was funded by the Mobility EU-Funds project CZ.02.2.69/0.0/0.0/16_027/0008503, awarded to the Crop Research Institute. Finishing of the work was supported by the institutional support from the Ministry of Agriculture of the Czech Republic - MZE-RO0418.

## Authors’ contribution

PS, WvdW and DB conceived the study; PS, DM and WvdW conceptualized the analysis; PS performed the literature search, extracted the data, performed the analysis, created tables and figures, and led writing of the manuscript; DM, DB and WvdW contributed to writing and edited the final version; all authors have read and approved the final version of this paper.

## Data availability

Data is available with the first author upon reasonable request.

