## Supplementary material for "Standardizing carabid pitfall catches for trapping effort; a meta-analysis": SupplemmentaryMaterials

List of appendices:

Appendix S1 **Literature search and article selection**

Appendix S2 **List of publications used for meta-analysis**

Appendix S3 **Frequency of studies and associated records according to data aggregation**

Appendix S4 **Overview of models fitted in this study**

Appendix S5 **Variation in continuous input variables related to trapping effort and pitfall catch**

Appendix S6 **Overview of model fits**

Appendix S7 **Comparing the subsets of data**

Appendix S8 **List of top models for model averaging**

Appendix S1

### Literature search and article selection

Three literature searches were made (Figure S1a). Two searches were conducted using as a database the Science Citation Index Expanded (SCI-E) through Web of Science (Clarivate). Search #1 covered the years of 1991-2018 (Figure S1a). Search #2 covered the years 1945-1990 and was less restrictive, because abstracts were not available for articles before 1991 in SCI-E (Figure S1a). Search #3 was made using Scopus (Elsevier) and covered the years 1960-2018 (Figure S1a). All searches were last updated on December 10, 2018. From the results of each search, reviews and studies outside the delimited geographical area (see below) were excluded. The results of the three searches were combined in EndNote v. 7. Duplicated publications were removed. The resulting pool of papers consisted of 648 publications (Figure S1b).

Article selection was done in two steps: (i) title and abstract screening; and, (ii) full text assessment (Figure S1b). The inclusion criteria applied to select the papers were:

- Data were based on pitfall traps, and carabid activity densities, species richness or both were provided for the entire community.
- We only included studies from annual field crops. These undergo periodical disturbances caused by field and crop management.
- Variation in field and crop management practices was addressed. The possible practices may include tillage, fertilization, crop rotation, cover crops, intercropping, weed control measures, use of pesticides and adoption of GM crops. Studies that focused on comparing farming systems (typically, organic vs. conventional) were also included.
- A geographical restriction was set to North America and Europe (i.e. including Russia west of the Ural).

Altogether, 544 articles were excluded since they did not satisfy these criteria or data could not be extracted (Figure S1b). At the end, 104 papers were retained and these formed the pool of publications for data extraction. The source publications are referenced in Appendix S2.

Data (n records = 810) were extracted from the main body of the paper, from tables, figures (using PlotDigitizer v. 2.6.8; Huwaldt & Steinhorst 2015) or from the supplementary materials associated with the publications on journals’ web pages. Authors of the original publications were contacted, if necessary, to clarify the sampling design or data presentation.

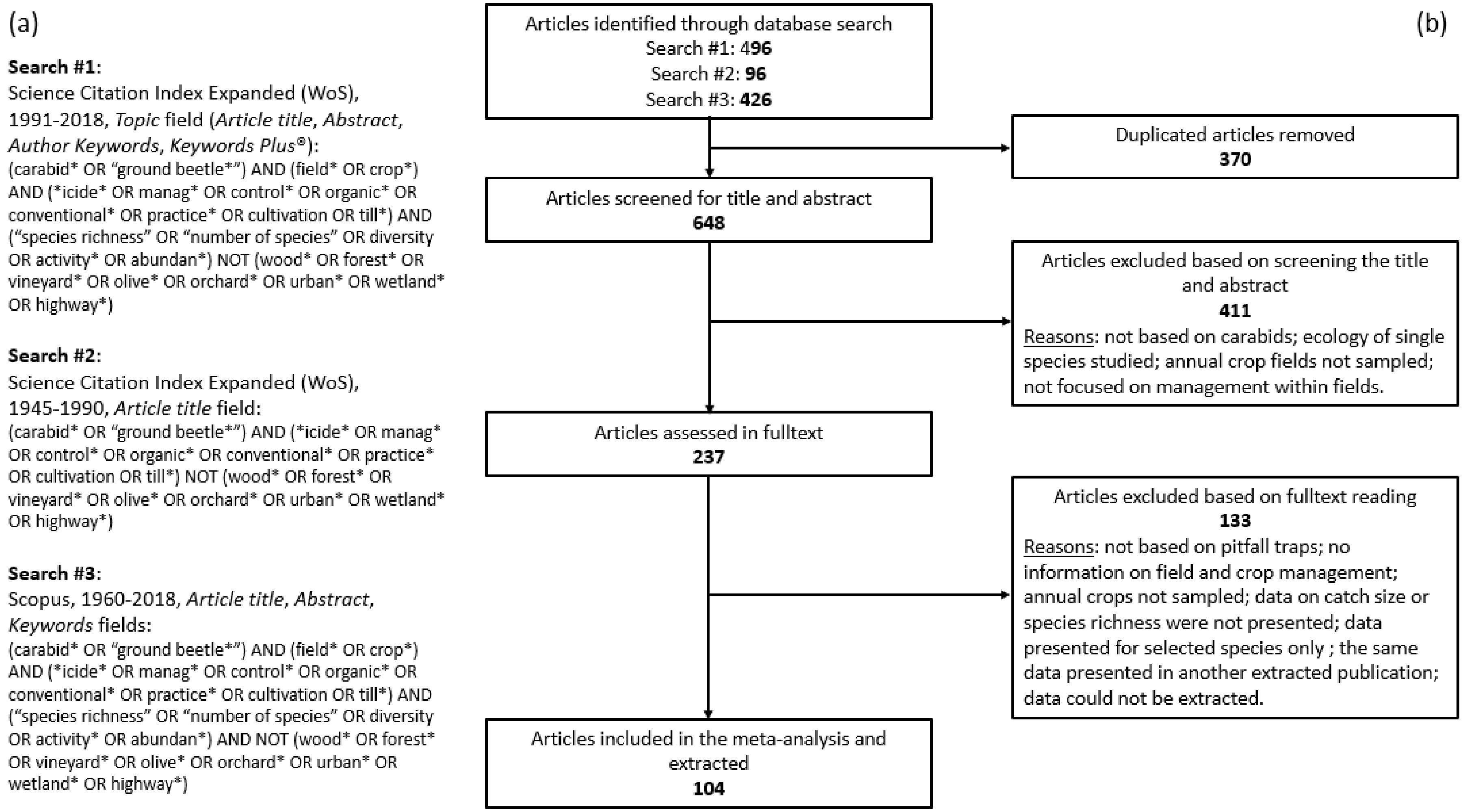


Figure S1. Description of literature search and article selection process. a) Search strings used to identify literature potentially useful for meta-analysis focused on the effect of field and crop management on carabid beetles. b) The PRISMA flow diagram showing the procedure used for selection of studies used in this study.

Appendix S2

### List of publications used for meta-analysis


Adhikari, S. & Menalled, F.D. (2018) Impacts of dryland farm management systems on weeds and ground beetles (carabidae) in the Northern Great Plains. Sustainability (Switzerland), 10, 2146.

Albajes, R., Farinos, G.P., Perez-Hedo, M., de la Poza, M., Lumbierres, B., Ortego, F., Pons, X. & Castanera, P. (2012) Post-market environmental monitoring of Bt maize in Spain: Non-target effects of varieties derived from the event MON810 on predatory fauna. Spanish Journal of Agricultural Research, 10, 977-985.

Albajes, R., Lopez, C. & Pons, X. (2003) Predatory fauna in cornfields and response to imidacloprid seed treatment. Journal of Economic Entomology, 96, 1805-1813.

Albajes, R., Lumbierres, B. & Pons, X. (2009) Responsiveness of Arthropod Herbivores and Their Natural Enemies to Modified Weed Management in Corn. Environmental Entomology, 38, 944-954.

Andersen, A. (1999) Plant protection in spring cereal production with reduced tillage. II. Pests and beneficial insects. Crop Protection, 18, 651-657.

Andersen, A. (2003) Long-term experiments with reduced tillage in spring cereals. II. Effects on pests and beneficial insects. Crop Protection, 22, 147-152.

Armstrong, G. (1995) Carabid beetle (Coleoptera, Carabidae) diversity and abundance in organic potatoes and conventionally grown seed potatoes in the north of Scotland. Pedobiologia, 39, 231-237.

Basedow, T. (2002) Changes in agriculture in an area in Northern Germany between the years 1971 and 2000, and the reactions of populations of predatory carabids (Col., Carabidae), of other predators, and of cereal aphids, to these changes. Zeitschrift für Pflanzenkrankheiten und Pflanzenschutz-Journal of Plant Diseases and Protection, 109, 1-14.

Basedow, T., Borg, A. & Scherney, F. (1976) Effects on insecticides upon terrestrial predaceous arthropods in cereal fields, especially ground beetles (Coleoptera, Carabidae). Entomologia Experimentalis et Applicata, 19, 37-51.

Batary, P., Holzschuh, A., Orci, K.M., Samu, F. & Tscharntke, T. (2012) Responses of plant, insect and spider biodiversity to local and landscape scale management intensity in cereal crops and grasslands. Agriculture, Ecosystems & Environment, 146, 130-136.

Belaoussoff, S., Kevan, P.G., Murphy, S. & Swanton, C. (2003) Assessing tillage disturbance on assemblages of ground beetles (Coleoptera: Carabidae) by using a range of ecological indices. Biodiversity and Conservation, 12, 851-882.

Bhatti, M.A., Duan, J., Head, G., Jiang, C.J., McKee, M.J., Nickson, T.E., Pilcher, C.L. & Pilcher, C.D. (2005) Field evaluation of the impact of corn rootworm (Coleoptera : Chrysomelidae)-protected Bt corn on ground-dwelling invertebrates. Environmental Entomology, 34, 1325-1335.

Boetel, M.A., Fuller, B.W., Chandler, L.D., Tollefson, J.J., McManus, B.L., Kadakia, N.D., Evenson, P.D. & Mishra, T.P. (2005) Nontarget arthropod abundance in areawide-managed corn habitats treated with semiochemical-based bait insecticide for corn rootworm (Coleoptera : Chrysomelidae) control. Journal of Economic Entomology, 98, 1957-1968.

Booij, C.J.H. & Noorlander, J. (1992) Farming systems and insect predators. Agriculture, Ecosystems and Environment, 40, 125-135.

Bourassa, S., Carcamo, H.A., Larney, F.J. & Spence, J.R. (2008) Carabid Assemblages (Coleoptera: Carabidae) in a Rotation of Three Different Crops in Southern Alberta, Canada: A Comparison of Sustainable and Conventional Farming. Environmental Entomology, 37, 1214-1223.

Bourassa, S., Carcamo, H.A., Spence, J.R., Blackshaw, R.E. & Floate, K. (2010) Effects of crop rotation and genetically modified herbicide-tolerant corn on ground beetle diversity, community structure, and activity density. Canadian Entomologist, 142, 143-159.

Burgio, G., Campanelli, G., Leteo, F., Ramilli, F., Depalo, L., Fabbri, R. & Sgolastra, F. (2015) Ecological Sustainability of an Organic Four-Year Vegetable Rotation System: Carabids and Other Soil Arthropods as Bioindicators. Agroecology and Sustainable Food Systems, 39, 295-316.

Butts, R.A., Floate, K.D., David, M., Blackshaw, R.E. & Burnett, P.A. (2003) Influence of intercropping canola or pea with barley on assemblages of ground beetles (Coleoptera : Carabidae). Environmental Entomology, 32, 535-541.

Carcamo, H.A. (1995) Effects of tillage on ground beetles (Coleoptera, Carabidae) – a farm-scale study in central Alberta. Canadian Entomologist, 127, 631-639.

Carcamo, H.A., Niemala, J.K. & Spence, J.R. (1995) Farming and ground beetles – effects of agronomic practice on populations and community structure. Canadian Entomologist, 127, 123-140.

Clark, M.S. (1999) Ground beetle abundance and community composition in conventional and organic tomato systems of California's Central Valley. Applied Soil Ecology, 11, 199-206.

Clark, M.S., Luna, J.M., Stone, N.D. & Youngman, R.R. (1993) Habitat preferences of generalist predators in reduced-tillage corn. Journal of Entomological Science, 28, 404-416.

Clark, S., Szlavecz, K., Cavigelli, M.A. & Purrington, F. (2006) Ground beetle (Coleoptera: Carabidae) assemblages in organic, no-till, and chisel-till cropping systems in Maryland. Environmental Entomology, 35, 1304-1312.

Clough, Y., Holzschuh, A., Gabriel, D., Purtauf, T., Kleijn, D., Kruess, A., Steffan-Dewenter, I. & Tscharntke, T. (2007) Alpha and beta diversity of arthropods and plants in organically and conventionally managed wheat fields. Journal of Applied Ecology, 44, 804-812.

Depalo, L., Burgio, G., von Fragstein, P., Kristensen, H.L., Bavec, M., Robacer, M., Campanelli, G. & Canali, S. (2017) Impact of living mulch on arthropod fauna: analysis of pest and beneficial dynamics on organic cauliflower (Brassica oleracea L. var. botrytis) in different European scenarios. Renewable Agriculture and Food Systems, 32, 240-247.

Diekotter, T., Wamser, S., Wolters, V. & Birkhofer, K. (2010) Landscape and management effects on structure and function of soil arthropod communities in winter wheat. Agriculture, Ecosystems & Environment, 137, 108-112.

Djoudi, E., Marie, A., Mangenot, A., Puech, C., Aviron, S., Plantegenest, M. & Petillon, J. (2018) Farming system and landscape characteristics differentially affect two dominant taxa of predatory arthropods. Agriculture, Ecosystems & Environment, 259, 98-110.

Djoudi, E.A., Plantegenest, M., Aviron, S. & Pétillon, J. (2019) Local vs. landscape characteristics differentially shape emerging and circulating assemblages of carabid beetles in agroecosystems. Agriculture, Ecosystems and Environment, 270-271, 149-158.

Dritschilo, W. & Erwin, T.L. (1982) Responses in abundance and diversity of cornfield carabid communities to differences in farm practices. Ecology, 63, 900-904.

Duan, J.J., Head, G., Jensen, A. & Reed, G. (2004) Effects of transgenic Bacillus thuringiensis potato and conventional insecticides for Colorado potato beetle (Coleoptera: Chrysomelidae) management on the abundance of ground-dwelling arthropods in Oregon potato ecosystems. Environmental Entomology, 33, 275-281.

Dunbar, M.W., Gassmann, A.J. & O'Neal, M.E. (2016) Impacts of Rotation Schemes on Ground-Dwelling Beneficial Arthropods. Environmental Entomology, 45, 1154-1160.

Dunbar, M.W., Gassmann, A.J. & O'Neal, M.E. (2017) Limited Impact of a Fall-Seeded, Spring-Terminated Rye Cover Crop on Beneficial Arthropods. Environmental Entomology, 46, 284-290.

Ekroos, J., Hyvonen, T., Tiainen, J. & Tiira, M. (2010) Responses in plant and carabid communities to farming practises in boreal landscapes. Agriculture, Ecosystems & Environment, 135, 288-293.

Ellsbury, M.M., Powell, J.E., Forcella, F., Woodson, W.D., Clay, S.A. & Riedell, W.E. (1998) Diversity and dominant species of ground beetle assemblages (Coleoptera: Carabidae) in crop rotation and chemical input systems for the northern Great Plains. Annals of the Entomological Society of America, 91, 619-625.

Eyre, M.D., Luff, M.L. & Leifert, C. (2013) Crop, field boundary, productivity and disturbance influences on ground beetles (Coleoptera, Carabidae) in the agroecosystem. Agriculture, Ecosystems & Environment, 165, 60-67.

Fan, Y.Q., Liebman, M., Groden, E. & Alford, A.R. (1993) Abundance of carabid beetles and other ground-dwelling arthropods in conventional versus low-input bean cropping systems. Agriculture, Ecosystems & Environment, 43, 127-139.

Farinos, G.P., de la Poza, M., Hernandez-Crespo, P., Ortego, F. & Castanera, P. (2008) Diversity and seasonal phenology of aboveground arthropods in conventional and transgenic maize crops in Central Spain. Biological Control, 44, 362-371.

Feber, R.E., Johnson, P.J., Bell, J.R., Chamberlain, D.E., Firbank, L.G., Fuller, R.J., Manley, W., Mathews, F., Norton, L.R., Townsend, M. & Macdonald, D.W. (2015) Organic Farming: Biodiversity Impacts Can Depend on Dispersal Characteristics and Landscape Context. Plos One, 10, e0135921.

Gailis, J. & Turka, I. (2014) The diversity and structure of ground beetles (Coleoptera: Carabidae) assemblages in differently managed winter wheat fields. Baltic Journal of Coleopterology, 14, 33-46.

Gallé, R., Happe, A.K., Baillod, A.B., Tscharntke, T. & Batáry, P. (2019) Landscape configuration, organic management, and within-field position drive functional diversity of spiders and carabids. Journal of Applied Ecology, 56, 63-72.

Garcia-Ruiz, E., Loureiro, I., Farinos, G.P., Gomez, P., Gutierrez, E., Sanchez, F.J., Escorial, M.C., Ortego, F., Chueca, M.C. & Castanera, P. (2018) Weeds and ground-dwelling predators' response to two different weed management systems in glyphosate-tolerant cotton: A farm-scale study. Plos One, 13, e0191408.

Groeneveld, J.H. & Klein, A.M. (2015) Pennycress-corn double-cropping increases ground beetle diversity. Biomass & Bioenergy, 77, 16-25.

Habustova, O.S., Svobodova, Z., Spitzer, L., Dolezal, P., Hussein, H.M. & Sehnal, F. (2015) Communities of ground-dwelling arthropods in conventional and transgenic maize: background data for the post-market environmental monitoring. Journal of Applied Entomology, 139, 31-45.

Hatten, T.D., Bosque-Perez, N.A., Labonte, J.R., Guy, S.O. & Eigenbrode, S.D. (2007) Effects of tillage on the activity density and biological diversity of carabid beetles in spring and winter crops. Environmental Entomology, 36, 356-368.

Helenius, J. (1990) Conventional and organic cropping systems at Suitia. 6. Insect populations in barley. Journal of Agricultural Science in Finland, 62, 349-355.

Hokkanen, H. & Holopainen, J.K. (1986) Carabid species and activity densities in biologically and conventionally managed cabbage fields. Journal of Applied Entomology-Zeitschrift Fur Angewandte Entomologie, 102, 353-363.

Hummel, J.D., Dosdall, L.M., Clayton, G.W., Harker, K.N. & O'Donovan, J.T. (2012) Ground Beetle (Coleoptera: Carabidae) Diversity, Activity Density, and Community Structure in a Diversified Agroecosystem. Environmental Entomology, 41, 72-80.

Irmler, U. (2003) The spatial and temporal pattern of carabid beetles on arable fields in northern Germany (Schleswig-Holstein) and their value as ecological indicators. Agriculture, Ecosystems & Environment, 98, 141-151.

Jabbour, R., Pisani-Gareau, T., Smith, R.G., Mullen, C. & Barbercheck, M. (2016) Cover crop and tillage intensities alter ground-dwelling arthropod communities during the transition to organic production. Renewable Agriculture and Food Systems, 31, 361-374.

Kalushkov, P., Gueorguiev, B., Spitzer, L. & Nedved, O. (2009) Biodiversity of ground beetles (Coleoptera: Carabidae) in genetically modified (Bt) and conventional (non-Bt) potato fields in Bulgaria. Biotechnology & Biotechnological Equipment, 23, 1346-1350.

Kennedy, P.J., Conrad, K.F., Perry, J.N., Powell, D., Aegerter, J., Todd, A.D., Walters, K.F.A. & Powell, W. (2001) Comparison of two field-scale approaches for the study of effects of insecticides on polyphagous predators in cereals. Applied Soil Ecology, 17, 253-266.

Kennedy, T.F., Connery, J., Fortune, T., Forristal, D. & Grant, J. (2013) A comparison of the effects of minimum-till and conventional-till methods, with and without straw incorporation, on slugs, slug damage, earthworms and carabid beetles in autumn-sown cereals. Journal of Agricultural Science, 151, 605-629.

Kocourek, F., Saska, P. & Rezac, M. (2013) Diversity of Carabid Beetles (Coleoptera: Carabidae) under Three Different Control Strategies against European Corn Borer in Maize. Plant Protection Science, 49, 146-153.

Kosewska, A. (2016) Conventional and non-inversion tillage systems as a factor causing changes in ground beetle (Col. Carabidae) assemblages in oilseed rape (Brassica napus) fields. Periodicum Biologorum, 118, 231-239.

Kosewska, A., Nietupski, M., Nijak, K. & Skalski, T. (2016) Effect of plant protection on assemblages of ground beetles (Coleoptera, Carabidae) in pea (Pisum L.) and lupine (Lupinus L.) crops. Periodicum Biologorum, 118, 213-222.

Kosewska, A., Skalski, T. & Nietupski, M. (2014) Effect of conventional and non-inversion tillage systems on the abundance and some life history traits of carabid beetles (Coleoptera: Carabidae) in winter triticale fields. European Journal of Entomology, 111, 669-676.

Kragten, S., Tamis, W.L.M., Gertenaar, E., Ramiro, S.M.M., Van der Poll, R.J., Wang, J. & De Snoo, G.R. (2011) Abundance of invertebrate prey for birds on organic and conventional arable farms in the Netherlands. Bird Conservation International, 21, 1-11.

Kromp, B. (1989) Carabid beetle communities (Carabidae, Coleoptera) in biologically and conventionally farmed agroecosystems. Agriculture, Ecosystems & Environment, 27, 241-251.

Kromp, B. (1990) Carabid beetles (Coleoptera, Carabidae) as bioindicators in biological conventional farming in Austrian potatoes fields. Biology and Fertility of Soils, 9, 182-187.

Kulagowski, R., Riggi, L. & Chailleux, A. (2016) Short-Term Effects of Conversion to Direct Seeding Mulch-Based Cropping Systems on Macro-Fauna and Weed Dynamics. Journal of Crop Improvement, 30, 65-83.

Lalonde, O., Legere, A., Stevenson, F.C., Roy, M. & Vanasse, A. (2012) Carabid beetle communities after 18 years of conservation tillage and crop rotation in a cool humid climate. Canadian Entomologist, 144, 645-657.

Langmaack, M., Land, S. & Buchs, W. (2001) Effects of different field management systems on the carabid coenosis in oil seed rape with special respect to ecology and nutritional status of predacious Poecilus cuprous L. (Col., Carabidae). Journal of Applied Entomology-Zeitschrift Fur Angewandte Entomologie, 125, 313-320.

Law, J.J. & Gallagher, R.S. (2018) Seed Distribution and Invertebrate Seed Predation in No-Till and Minimum-Till Maize Systems. Agronomy Journal, 110, 2488-2495.

Leslie, T.W., Biddinger, D.J., Rohr, J.R. & Fleischer, S.J. (2010) Conventional and Seed-Based Insect Management Strategies Similarly Influence Nontarget Coleopteran Communities in Maize. Environmental Entomology, 39, 2045-2055.

Leslie, T.W., Hoheisel, G.A., Biddinger, D.J., Rohr, J.R. & Fleischer, S.J. (2007) Transgenes sustain epigeal insect biodiversity in diversified vegetable farm systems. Environmental Entomology, 36, 234-244.

Lewis, M.T., Fleischer, S.J. & Roberts, D.C. (2016) Horticultural Production Systems Influence Ground Beetle (Coleoptera: Carabidae) Distribution and Diversity in Cucurbits. Environmental Entomology, 45, 559-569.

Lopez, M.D., Prasifka, J.R., Bruck, D.J. & Lewis, L.C. (2005) Utility of Ground Beetle Species in Field Tests of Potential Nontarget Effects of Bt Crops. Environmental Entomology, 34, 1317-1324.

MacDonald, M.A., Cobbold, G., Mathews, F., Denny, M.J.H., Walker, L.K., Grice, P.V. & Anderson, G.Q.A. (2012) Effects of agri-environment management for cirl buntings on other biodiversity. Biodiversity and Conservation, 21, 1477-1492.

Magagnoli, S., Masetti, A., Depalo, L., Sommaggio, D., Campanelli, G., Leteo, F., Lovei, G.L. & Burgio, G. (2018) Cover crop termination techniques affect ground predation within an organic vegetable rotation system: A test with artificial caterpillars. Biological Control, 117, 109-114.

Menalled, F.D., Smith, R.G., Dauer, J.T. & Fox, T.B. (2007) Impact of agricultural management on carabid communities and weed seed predation. Agriculture, Ecosystems & Environment, 118, 49-54.

O'Rourke, M.E., Liebman, M. & Rice, M.E. (2008) Ground beetle (Coleoptera: Carabidae) assemblages in conventional and diversified crop rotation systems. Environmental Entomology, 37, 121-130.

O'Sullivan, C.M. & Gormally, M.J. (2002) A comparison of ground beetle (Carabidae: Coleoptera) communities in an organic and conventional potato crop. Biological Agriculture & Horticulture, 20, 99-110.

Pavuk, D.M., Purrington, F.F., Williams, C.E. & Stinner, B.R. (1997) Ground beetle (Coleoptera: Carabidae) activity density and community composition in vegetationally diverse corn agroecosystems. American Midland Naturalist, 138, 14-28.

Pfiffner, L. & Luka, H. (2003) Effects of low-input farming systems on carabids and epigeal spiders - a paired farm approach. Basic and Applied Ecology, 4, 117-127.

Pfiffner, L. & Niggli, U. (1996) Effects of bio-dynamic, organic and conventional farming on ground beetles (Col Carabidae) and other epigaeic arthropods in winter wheat. Biological Agriculture & Horticulture, 12, 353-364.

Pocock, M.J.O. & Jennings, N. (2008) Testing biotic indicator taxa: the sensitivity of insectivorous mammals and their prey to the intensification of lowland agriculture. Journal of Applied Ecology, 45, 151-160.

Ponce, C., Bravo, C., de León, D.G., Magaña, M. & Alonso, J.C. (2011) Effects of organic farming on plant and arthropod communities: A case study in Mediterranean dryland cereal. Agriculture, Ecosystems & Environment, 141, 193-201.

Porhajašová, J., Petřvalský, V., Šustek, Z., Urminská, J., Ondrišík, P. & Noskovič, J. (2008) Long-termed changes in ground beetle (Coleoptera: Carabidae) assemblages in a field treated by organic fertilizers. Biologia, 63, 1184-1195.

Prasifka, J.R., Schmidt, N.P., Kohler, K.A., O'Neal, M.E., Hellmich, R.L. & Singer, J.W. (2006) Effects of living mulches on predator abundance and sentinel prey in a corn-soybean-forage rotation. Environmental Entomology, 35, 1423-1431.

Puech, C., Poggi, S., Baudry, J. & Aviron, S. (2015) Do farming practices affect natural enemies at the landscape scale? Landscape Ecology, 30, 125-140.

Purtauf, T., Roschewitz, I., Dauber, J., Thies, C., Tscharntke, T. & Wolters, V. (2005) Landscape context of organic and conventional farms: Influences on carabid beetle diversity. Agriculture, Ecosystems & Environment, 108, 165-174.

Rivers, A.N., Mullen, C.A. & Barbercheck, M.E. (2018) Cover Crop Species and Management Influence Predatory Arthropods and Predation in an Organically Managed, Reduced-Tillage Cropping System. Environmental Entomology, 47, 340-355.

Seagraves, M.P., McPherson, R.M. & Ruberson, J.R. (2004) Impact of Solenopsis invicta Buren suppression on arthropod ground predators and pest species in soybean. Journal of Entomological Science, 39, 433-443.

Sereda, E., Wolters, V. & Birkhofer, K. (2015) Addition of crop residues affects a detritus-based food chain depending on litter type and farming system. Basic and Applied Ecology, 16, 746-754.

Shah, P.A., Brooks, D.R., Ashby, J.E., Perry, J.N. & Woiwod, I.P. (2003) Diversity and abundance of the coleopteran fauna from organic and conventional management systems in southern England. Agricultural and Forest Entomology, 5, 51-60.

Schier, A. (2006) Field study on the occurrence of ground beetles and spiders in genetically modified, herbicide tolerant corn in conventional and conservation tillage systems. Journal of Plant Diseases and Protection, 113 (Special Issue), S101-S113.

Schorling, M. & Freier, B. (2006) Six-year monitoring of non-target arthropods in Bt maize (Cry 1Ab) in the European corn borer (Ostrinia nubilalis) infestation area Oderbruch (Germany). Journal fur Verbraucherschutz und Lebensmittelsicherheit, 1, 106-108.

Staudacher, K., Rubbmark, O.R., Birkhofer, K., Malsher, G., Sint, D., Jonsson, M. & Traugott, M. (2018) Habitat heterogeneity induces rapid changes in the feeding behaviour of generalist arthropod predators. Functional Ecology, 32, 809-819.

Stephens, E.J., Losey, J.E., Allee, L.L., DiTommaso, A., Bodner, C. & Breyre, A. (2012) The impact of Cry3Bb Bt-maize on two guilds of beneficial beetles. Agriculture, Ecosystems & Environment, 156, 72-81.

Svobodova, Z., Habustova, O.S., Holec, J., Holec, M., Bohac, J., Jursik, M., Soukup, J. & Sehnal, F. (2018) Split application of glyphosate in herbicide-tolerant maize provides efficient weed control and favors beneficial epigeic arthropods. Agriculture, Ecosystems & Environment, 251, 171-179.

Tamutis, V., Monsevičius, V. & Pekarskas, J. (2004) Ground and rove beetles (Coleoptera: Carabidae, Staphylinidae) in ecological and conventional winter wheat fields. Baltic Journal of Coleopterology, 4, 31-40.

Trichard, A., Ricci, B., Ducourtieux, C. & Petit, S. (2014) The spatio-temporal distribution of weed seed predation differs between conservation agriculture and conventional tillage. Agriculture, Ecosystems & Environment, 188, 40-47.

Twardowski, J.P., Beres, P., Hurej, M., Klukowski, Z. & Warzecha, R. (2017) Effects of maize expressing the insecticidal protein Cry1AB on non-target ground beetle assemblages (Coleoptera, Carabidae). Romanian Agricultural Research, 34, 351-361.

van der Laat, R., Owen, M.D.K., Liebman, M. & Leon, R.G. (2015) Postdispersal Weed Seed Predation and Invertebrate Activity Density in Three Tillage Regimes. Weed Science, 63, 828-838.

Vesely, M. & Sarapatka, B. (2008) Effects of conversion to organic farming on carabid beetles (Carabidae) in experimental fields in the Czech Republic. Biological Agriculture & Horticulture, 25, 289-309.

Vician, V., Svitok, M., Kočík, K. & Stašiov, S. (2015) The influence of agricultural management on the structure of ground beetle (Coleoptera: Carabidae) assemblages. Biologia (Poland), 70, 240-251.

Volkmar, C., Hussein, M.L.A. & Wetzel, T. (2004) Ecological field studies in transgenic maize at Friemar (Thuringia). Zeitschrift für Pflanzenkrankheiten und Pflanzenschutz-Journal of Plant Diseases and Protection, 111 (Special Issue), S1017-S1024.

Volkmar, C., Kreuter, T., Richter, L., Lubke-Al Hussein, M., Jany, D., Schmutzler, K. & Wetzel, T. (2000) Ecological studies accompanying the cultivation of transgenic and conventional rape plants in the Central German region from 1996 to 1998. Zeitschrift für Pflanzenkrankheiten und Pflanzenschutz-Journal of Plant Diseases and Protection, 107 (Special Issue), S337-S345.

Volkmar, C., Lübke-Al Hussein, M. & Kreuter, T. (2003) Effects of reduced soil tillage on the activity of epigeic arthropods. Gesunde Pflanzen, 55, 40-45.

Volkmar, C., Lubke-Al Hussein, M., Kreuter, T. & Wetzel, T. (2002) Does the cultivation of transgenic sugar beet affect the stability of arthropod coenoses (results of a four-year field study)? Zeitschrift für Pflanzenkrankheiten und Pflanzenschutz-Journal of Plant Diseases and Protection, 109 (Special Issue), S1031-S1038.

Volkmar, C. & Schützel, A. (2008) Ecological study of farming sites in Saxony-Anhalt and possibilities of its application in monitoring and subsidy programmes. Archives of Phytopathology and Plant Protection, 41, 129-141.

Wick, M. & Freier, B. (2000) Long-term effects of an insecticide application on non-target arthropods in winter wheat - a field study over two seasons. Anzeiger Fur Schadlingskunde-Journal of Pest Science, 73, 61-69.

Wick, M., Freier, B., Kreuter, T. & Moll, E. (2001) Effects of insecticide application to winter wheat and subsequent tillage on groundbeetle communities (Coleoptera: Carabidae). Entomologia Generalis, 25, 265-273.

Witmer, J.E., Hough-Goldstein, J.A. & Pesek, J.D. (2003) Ground-dwelling and foliar arthropods in four cropping systems. Environmental Entomology, 32, 366-376.

Appendix S3

**Frequency of studies and associated records according to data aggregation**

Table S1 Number of studies and number of data records for each of six types of study design

| Type of experiment | Level of data aggregation in source publications | | Number of studies | Number of records |
|---|---|---|---|---|
| | Reporting of data over replicates | Reporting of data over seasons | | |
| Field experiments with replicate plots | Means or totals per treatment | Per each season | 50 | 536 |
| | | Over multiple seasons | 13 | 68 |
| Whole field experiments | Means or totals per treatment | Per each season | 19 | 78 |
| | | Over multiple seasons | 4 | 12 |
| | Observations in each field | Per each season | 16 | 112 |
| | | Over multiple seasons | 2 | 4 |

Appendix S4

**Overview of models fitted in this study**

Table S1. List of fitted models that were compared in order to find the optimal standardization of the pitfall catch for unit effort of trapping. The random and error terms are assumed to be normally distributed. Weights $w$ in model C27 were defined as log($R$), and in model S27 as log($Q^{0.25}$).

$C$ – number of individuals caught; $S$ – observed species richness; $K$ – number traps used; $P$ – total perimeter of traps used; $X$ – duration of trap exposure; $R$ – trap-days, i.e. $XK$; $Q$ – perimeter-days, i.e. $XP$; $i$ – independent study; $j$ – year of sampling; $k$ – record; $a_i$ or $a_{0i}$ – random intercept for a study $i$; $b_{ij}$ or $b_{0ij}$ – random intercept for the year of sampling $j$ nested within a study $i$; $a_{1i}$ – random slope for a study $i$; $b_{1ij}$ – random slope for the year of sampling $j$ nested within a study $i$; $\epsilon$ – error.

| # | Total catch | $w$ |
|---|---|---|
| C1 | $\log(C_k) = \beta_0 + \beta_1\log(K_k) + \epsilon_k$ | No |
| C2 | $\log(C_{ik}) = \beta_0 + \beta_1\log(K_{ik}) + a_i + \epsilon_{ik}$ | No |
| C3 | $\log(C_{ijk}) = \beta_0 + \beta_1\log(K_{ijk}) + a_i + b_{ij} + \epsilon_{ijk}$ | No |
| C4 | $\log(C_{ik}) = (\beta_0 + a_{0i}) + (\beta_1 + a_{1i})\log(K_{ik}) + \epsilon_{ik}$ | No |
| C5 | $\log(C_{ijk}) = (\beta_0 + a_{0i} + b_{0ij}) + (\beta_1 + a_{1i} + b_{1ij})\log(K_{ijk}) + \epsilon_{ijk}$ | No |
| C6 | $\log(C_k) = \beta_0 + \beta_1\log(P_k) + \epsilon_k$ | No |
| C7 | $\log(C_{ik}) = \beta_0 + \beta_1\log(P_{ik}) + a_i + \epsilon_{ik}$ | No |
| C8 | $\log(C_{ijk}) = \beta_0 + \beta_1\log(P_{ijk}) + a_i + b_{ij} + \epsilon_{ijk}$ | No |
| C9 | $\log(C_{ik}) = (\beta_0 + a_{0i}) + (\beta_1 + a_{1i})\log(P_{ik}) + \epsilon_{ik}$ | No |
| C10 | $\log(C_{ijk}) = (\beta_0 + a_{0i} + b_{0ij}) + (\beta_1 + a_{1i} + b_{1ij})\log(P_{ijk}) + \epsilon_{ijk}$ | No |
| C11 | $\log(C_k) = \beta_0 + \beta_1\log(X_k) + \epsilon_k$ | No |
| C12 | $\log(C_{ik}) = \beta_0 + \beta_1\log(X_{ik}) + a_i + \epsilon_{ik}$ | No |
| C13 | $\log(C_{ijk}) = \beta_0 + \beta_1\log(X_{ijk}) + a_i + b_{ij} + \epsilon_{ijk}$ | No |
| C14 | $\log(C_{ik}) = (\beta_0 + a_{0i}) + (\beta_1 + a_{1i})\log(X_{ik}) + \epsilon_{ik}$ | No |
| C15 | $\log(C_{ijk}) = (\beta_0 + a_{0i} + b_{0ij}) + (\beta_1 + a_{1i} + b_{1ij})\log(X_{ijk}) + \epsilon_{ijk}$ | No |
| C16 | $\log(C_k) = \beta_0 + \beta_1\log(R_k) + \epsilon_k$ | No |
| C17 | $\log(C_{ik}) = \beta_0 + \beta_1\log(R_{ik}) + a_i + \epsilon_{ik}$ | No |
| C18 | $\log(C_{ijk}) = \beta_0 + \beta_1\log(R_{ijk}) + a_i + b_{ij} + \epsilon_{ijk}$ | No |
| C19 | $\log(C_{ik}) = (\beta_0 + a_{0i}) + (\beta_1 + a_{1i})\log(R_{ik}) + \epsilon_{ik}$ | No |
| C20 | $\log(C_{ijk}) = (\beta_0 + a_{0i} + b_{0ij}) + (\beta_1 + a_{1i} + b_{1ij})\log(R_{ijk}) + \epsilon_{ijk}$ | No |
| C21 | $\log(C_k) = \beta_0 + \beta_1\log(Q_k) + \epsilon_k$ | No |
| C22 | $\log(C_{ik}) = \beta_0 + \beta_1\log(Q_{ik}) + a_i + \epsilon_{ik}$ | No |
| C23 | $\log(C_{ijk}) = \beta_0 + \beta_1\log(Q_{ijk}) + a_i + b_{ij} + \epsilon_{ijk}$ | No |
| C23 | $\log(C_{ik}) = (\beta_0 + a_{0i}) + (\beta_1 + a_{1i})\log(Q_{ik}) + \epsilon_{ik}$ | No |
| C25 | $\log(C_{ijk}) = (\beta_0 + a_{0i} + b_{0ij}) + (\beta_1 + a_{1i} + b_{1ij})\log(Q_{ijk}) + \epsilon_{ijk}$ | No |
| C26 | $\log(C_{ijk}) = \beta_0 + \beta_1(\log(R_{ijk}) - \log(\bar{R})) + a_i + b_{ij} + \epsilon_{ijk}$ | No |
| C27 | $\log(C_{ijk}) = \beta_0 + \beta_1(\log(R_{ijk}) - \log(\bar{R})) + a_i + b_{ij} + \epsilon_{ijk}$ | Yes |

| | Species richness | |
|---|---|---|
| S1 | $S_k \sim Poisson(\lambda_k), \log(\lambda_k) = \beta_0 + \beta_1 \log(K_k)$ | No |
| S2 | $S_{ik} \sim Poisson(\lambda_{ik}), \log(\lambda_{ik}) = \beta_0 + \beta_1 \log(K_{ik}) + a_i$ | No |
| S3 | $S_{ijk} \sim Poisson(\lambda_{ijk}), \log(\lambda_{ijk}) = \beta_0 + \beta_1 \log(K_{ijk}) + a_i + b_{ij}$ | No |
| S4 | $S_{ik} \sim Poisson(\lambda_{ik}), \log(\lambda_{ik}) = (\beta_0 + a_{0i}) + (\beta_1 + a_{1i}) \log(K_{ik})$ | No |
| S5 | $S_{ijk} \sim Poisson(\lambda_{ijk}), \log(\lambda_{ijk}) = (\beta_0 + a_{0i} + b_{0ij}) + (\beta_1 + a_{1i} + b_{1ij}) \log(K_{ijk})$ | No |
| S6 | $S_k \sim Poisson(\lambda_k), \log(\lambda_k) = \beta_0 + \beta_1 \log(P_k)$ | No |
| S7 | $S_{ik} \sim Poisson(\lambda_{ik}), \log(\lambda_{ik}) = \beta_0 + \beta_1 \log(P_{ik}) + a_i$ | No |
| S8 | $S_{ijk} \sim Poisson(\lambda_{ijk}), \log(\lambda_{ijk}) = \beta_0 + \beta_1 \log(P_{ijk}) + a_i + b_{ij}$ | No |
| S9 | $S_{ik} \sim Poisson(\lambda_{ik}), \log(\lambda_{ik}) = (\beta_0 + a_{0i}) + (\beta_1 + a_{1i}) \log(P_{ik})$ | No |
| S10 | $S_{ijk} \sim Poisson(\lambda_{ijk}), \log(\lambda_{ijk}) = (\beta_0 + a_{0i} + b_{0ij}) + (\beta_1 + a_{1i} + b_{1ij}) \log(P_{ijk})$ | No |
| S11 | $S_k \sim Poisson(\lambda_k), \log(\lambda_k) = \beta_0 + \beta_1 \log(X_k)$ | No |
| S12 | $S_{ik} \sim Poisson(\lambda_{ik}), \log(\lambda_{ik}) = \beta_0 + \beta_1 \log(X_{ik}) + a_i$ | No |
| S13 | $S_{ijk} \sim Poisson(\lambda_{ijk}), \log(\lambda_{ijk}) = \beta_0 + \beta_1 \log(X_{ijk}) + a_i + b_{ij}$ | No |
| S14 | $S_{ik} \sim Poisson(\lambda_{ik}), \log(\lambda_{ik}) = (\beta_0 + a_{0i}) + (\beta_1 + a_{1i}) \log(X_{ik})$ | No |
| S15 | $S_{ijk} \sim Poisson(\lambda_{ijk}), \log(\lambda_{ijk}) = (\beta_0 + a_{0i} + b_{0ij}) + (\beta_1 + a_{1i} + b_{1ij}) \log(X_{ijk})$ | No |
| S16 | $S_k \sim Poisson(\lambda_k), \log(\lambda_k) = \beta_0 + \beta_1 \log(R_k)$ | No |
| S17 | $S_{ik} \sim Poisson(\lambda_{ik}), \log(\lambda_{ik}) = \beta_0 + \beta_1 \log(R_{ik}) + a_i$ | No |
| S18 | $S_{ijk} \sim Poisson(\lambda_{ijk}), \log(\lambda_{ijk}) = \beta_0 + \beta_1 \log(R_{ijk}) + a_i + b_{ij}$ | No |
| S19 | $S_{ik} \sim Poisson(\lambda_{ik}), \log(\lambda_{ik}) = (\beta_0 + a_{0i}) + (\beta_1 + a_{1i}) \log(R_{ik})$ | No |
| S20 | $S_{ijk} \sim Poisson(\lambda_{ijk}), \log(\lambda_{ijk}) = (\beta_0 + a_{0i} + b_{0ij}) + (\beta_1 + a_{1i} + b_{1ij}) \log(R_{ijk})$ | No |
| S21 | $S_k \sim Poisson(\lambda_k), \log(\lambda_k) = \beta_0 + \beta_1 \log(Q_k)$ | No |
| S22 | $S_{ik} \sim Poisson(\lambda_{ik}), \log(\lambda_{ik}) = \beta_0 + \beta_1 \log(Q_{ik}) + a_i$ | No |
| S23 | $S_{ijk} \sim Poisson(\lambda_{ijk}), \log(\lambda_{ijk}) = \beta_0 + \beta_1 \log(Q_{ijk}) + a_i + b_{ij}$ | No |
| S24 | $S_{ik} \sim Poisson(\lambda_{ik}), \log(\lambda_{ik}) = (\beta_0 + a_{0i}) + (\beta_1 + a_{1i}) \log(Q_{ik})$ | No |
| S25 | $S_{ijk} \sim Poisson(\lambda_{ijk}), \log(\lambda_{ijk}) = (\beta_0 + a_{0i} + b_{0ij}) + (\beta_1 + a_{1i} + b_{1ij}) \log(Q_{ijk})$ | No |
| S26 | $S_{ijk} \sim Poisson(\lambda_{ijk}), \log(\lambda_{ijk}) = \beta_0 + \beta_1 (\log(Q_{ijk}) - \log(\bar{Q})) + a_i + b_{ij}$ | No |
| S27 | $S_{ijk} \sim Poisson(\lambda_{ijk}), \log(\lambda_{ijk}) = \beta_0 + \beta_1 (\log(Q_{ijk}) - \log(\bar{Q})) + a_i + b_{ij}$ | Yes |

Table S2 List of models fitted to explore which variables fundamental to the design and level of data aggregation in source publications affect the total catch size per unit effort, and recorded species richness per unit effort. Variables *Continent*, *Unit*, *RowWidth*, *Season*, *Fluid*, *Funnel* are categorical, *Year* is continuous. The random and error terms are assumed to be normally distributed. Weights $w$ in models for $C$ were defined as log($R$).

$C$ – number of individuals caught; $S$ – observed species richness; $R$ – trap-days; $Q$ – perimeter-days; $i$ – independent study; $j$ – year of sampling; $k$ – record; $\xi_{ijk}$ – offset defined as $R_{ijk}$ for catch size and as $Q_{ijk}^{0.25}$ for species richness; $a_i$ – random intercept for a study $i$; $b_{ij}$ – random intercept for the year of sampling $j$ nested within a study $i$; $\epsilon$ – error.

| # | Total catch | $w$ |
|---|---|---|
| C28 | $\log(C_{ijk}/\xi_{ijk}) = \beta_0 + Continent_{ijk} + a_i + b_{ij} + \epsilon_{ijk}$ | Yes |
| C29 | $\log(C_{ijk}/\xi_{ijk}) = \beta_0 + Unit_{ijk} + a_i + b_{ij} + \epsilon_{ijk}$ | Yes |
| C30 | $\log(C_{ijk}/\xi_{ijk}) = \beta_0 + RowWidth_{ijk} + a_i + b_{ij} + \epsilon_{ijk}$ | Yes |
| C31 | $\log(C_{ijk}/\xi_{ijk}) = \beta_0 + Season_{ijk} + a_i + b_{ij} + \epsilon_{ijk}$ | Yes |
| C32 | $\log(C_{ijk}/\xi_{ijk}) = \beta_0 + Funnel_{ijk} + a_i + b_{ij} + \epsilon_{ijk}$ | Yes |
| C34 | $\log(C_{ijk}/\xi_{ijk}) = \beta_0 + Fluid_{ijk} + a_i + b_{ij} + \epsilon_{ijk}$ | Yes |
| C35 | $\log(C_{ijk}/\xi_{ijk}) = \beta_0 + \beta_1\, Year_{ijk} + a_i + b_{ij} + \epsilon_{ijk}$ | Yes |
| | Species richness | |
| S28 | $S_{ijk} \sim Poisson(\lambda_{ijk}/\xi_{ijk}), \log(\lambda_{ijk}/\xi_{ijk}) = \beta_0 + Continent_{ijk} + a_i + b_{ij}$ | No |
| S29 | $S_{ijk} \sim Poisson(\lambda_{ijk}/\xi_{ijk}), \log(\lambda_{ijk}/\xi_{ijk}) = \beta_0 + Unit_{ijk} + a_i + b_{ij}$ | No |
| S30 | $S_{ijk} \sim Poisson(\lambda_{ijk}/\xi_{ijk}), \log(\lambda_{ijk}/\xi_{ijk}) = \beta_0 + RowWidth_{ijk} + a_i + b_{ij}$ | No |
| S31 | $S_{ijk} \sim Poisson(\lambda_{ijk}/\xi_{ijk}), \log(\lambda_{ijk}/\xi_{ijk}) = \beta_0 + Season_{ijk} + a_i + b_{ij}$ | No |
| S32 | $S_{ijk} \sim Poisson(\lambda_{ijk}/\xi_{ijk}), \log(\lambda_{ijk}/\xi_{ijk}) = \beta_0 + Funnel_{ijk} + a_i + b_{ij}$ | No |
| S34 | $S_{ijk} \sim Poisson(\lambda_{ijk}/\xi_{ijk}), \log(\lambda_{ijk}/\xi_{ijk}) = \beta_0 + Fluid_{ijk} + a_i + b_{ij}$ | No |
| S35 | $S_{ijk} \sim Poisson(\lambda_{ijk}/\xi_{ijk}), \log(\lambda_{ijk}/\xi_{ijk}) = \beta_0 + \beta_1\, Year_{ijk} + a_i + b_{ij}$ | No |

Table S3. List of models that were fitted in order to explore the importance of variables fundamental to the design and level of data aggregation in source publications for noise they presumably systematically introduce to the relationships of the pitfall catch size with unit effort of pitfall trapping. Variables *Fluid*, *Funnel*, *Continent*, *Unit*, *RowWidth* and *Season* are categorical, *Year* is continuous. The random and error terms are assumed to be normally distributed. Weights *w* in models for *C* were defined as log(*R*).

*C* – number of individuals caught; *S* – observed species richness; *R* – trap-days; *Q* – perimeter-days; *i* – independent study; *j* – year of sampling; *k* – record; $\xi_{ijk}$ – offset defined as $R_{ijk}$ for catch size and as $Q_{ijk}^{0.25}$ for species richness; $a_i$ – random intercept for a study *i*; $b_{ij}$ – random intercept for the year of sampling *j* nested within a study *i*; $\epsilon$ – error;

| # | Total catch | *w* |
|---|---|---|
| C36 | $\log(I_{ijk}/\xi_{ijk}) = \gamma_0 + Continent_{ijk} + Unit_{ijk} + RowWidth_{ijk} + Season_{ijk} + Funnel_{ijk} + Fluid_{ijk} + \gamma_1 Year_{ijk} + a_i + b_{ij} + \epsilon_{ijk}$ | Yes |
| | Species richness | |
| S36 | $S_{ijk} \sim Poisson(\lambda_{ijk}/\xi_{ijk}), \log(\lambda_{ijk}/\xi_{ijk}) = \gamma_0 + Continent_{ijk} + Unit_{ijk} + RowWidth_{ijk} + Season_{ijk} + Funnel_{ijk} + Fluid_{ijk} + \gamma_1 Year_{ijk} + a_i + b_{ij}$ | No |

Appendix S5

**Variation in continuous input variables related to trapping effort and carabid catch.**

Table S1 Variation in continuous input variables related to trapping effort and carabid catch

$K$ – number of traps used; $d$ – perimeter of the circular traps used; $l$ – length of the square traps used; $P$ – total perimeter of the traps used calculated as $P = K\pi d$ for circular traps or as $P = 4Kl$ for quadratic traps; $X$ – exposure time of the traps; $R$ – trap-days calculated as $R = KX$; $Q$ – perimeter-days calculated as $Q = PX$; $I$ – number of individuals collected; $S$ – observed species richness.

| Variable | Unit | n records | mean | min | max | median |
|---|---|---|---|---|---|---|
| *K* | [trap] | 810 | 17.1 | 3 | 540 | 10 |
| *d* | [m] | 804 | 0.097 | 0.056 | 0.125 | 0.10 |
| *l* | [m] | 6 | 0.106 | | | |
| *P* | [m] | 810 | 5.0 | 0.8 | 118.8 | 3.1 |
| *X* | [day] | 810 | 74.1 | 2 | 420 | 58 |
| *R* | [trap day] | 810 | 944.3 | 20 | 15120 | 525 |
| *Q* | [m day] | 810 | 268.7 | 6.3 | 3845.3 | 159.4 |
| *C* | [individuals] | 792 | 1898.7 | 21 | 47424 | 652 |
| *S* | [species] | 335 | 27.1 | 5 | 69 | 26 |

Appendix S6

### Overview of the fitted models to find the best standardization

Table S1 Overview of model fits for catch per unit effort relationships for total catch (*C*) and species richness of the catch (*S*). Models differ in the measure of trapping effort (*K*: models C1-5 and S1-5; *P*: models C6-10 and S6-10; *t*: models C11-15 and S11-15; *R*: models C16-20 and S16-20; *Q*: models C21-25 and S21-25) and in the structure of the random effects (Appendix S4, Table 1). Akaike's Information Criterion (AIC) and Bayesian Information Criterion (BIC) were estimated using maximum likelihood. Slopes and their s.e. are given as estimated from the models. All slopes are significantly different from 0 at α<0.001. The best model has a ΔAIC or ΔBIC of zero. The preferred models are highlighted in bold.

| Total catch | | | | | Species richness | | | | |
|---|---|---|---|---|---|---|---|---|---|
| Model | ΔAIC | ΔBIC | $\beta_1$ | s.e. | Model | Δ*AIC* | Δ*BIC* | $\beta_1$ | s.e. |
| C1 | 1287.9 | 1268.6 | 0.462 | 0.06 | S1 | 964.2 | 956.6 | 0.303 | 0.012 |
| C2 | 333 | 318.4 | 0.638 | 0.091 | S2 | 62.1 | 58.3 | 0.281 | 0.048 |
| C3 | 151.9 | 141.9 | 0.726 | 0.082 | S3 | 13.7 | 13.7 | 0.284 | 0.05 |
| C4 | 329.4 | 324.1 | 0.566 | 0.123 | S4 | 65.8 | 69.6 | 0.289 | 0.054 |
| C5 [c] | 145.6 | 154.4 | 0.588 | 0.127 | S5 [s] | 18.6 | 33.9 | 0.292 | 0.059 |
| C6 | 1308.2 | 1288.9 | 0.375 | 0.06 | S6 | 973.1 | 965.5 | 0.303 | 0.012 |
| C7 | 335.3 | 320.7 | 0.629 | 0.092 | S7 | 52.8 | 49 | 0.319 | 0.05 |
| C8 | 157.8 | 147.8 | 0.706 | 0.083 | S8 | 11.9 | 11.9 | 0.295 | 0.051 |
| C9 | 326.4 | 321.2 | 0.464 | 0.148 | S9 | 42.6 | 46.4 | 0.324 | 0.087 |
| C10 [c] | 150.6 | 159.4 | 0.522 | 0.134 | S10 [s] | 17.1 | 32.3 | 0.302 | 0.059 |
| C11 | 1093.4 | 1074.1 | 0.723 | 0.042 | S11 | 1515.9 | 1508.2 | 0.06 | 0.01 |
| C12 | 247.9 | 233.3 | 0.893 | 0.074 | S12 | 78.5 | 74.7 | 0.168 | 0.046 |
| C13 | 140.7 | 130.7 | 0.837 | 0.086 | S13 | 36.7 | 36.7 | 0.131 | 0.053 |
| C14 [c] | 239 | 233.7 | 0.914 | 0.115 | S14 | 82.4 | 86.2 | 0.163 | 0.063 |
| C15 [c] | 144.2 | 153 | 0.833 | 0.098 | S15 [s] | 41.4 | 56.6 | 0.137 | 0.055 |
| C16 | 751 | 731.7 | 0.995 | 0.033 | S16 | 993.8 | 986.1 | 0.222 | 0.01 |
| C17 | 136.9 | 122.3 | 0.953 | 0.054 | S17 [c] | 42.7 | 38.9 | 0.247 | 0.033 |
| **C18** | **9.9** | **0** | **0.938** | **0.056** | S18 | 3.4 | 3.4 | 0.249 | 0.037 |
| C19 | 114.1 | 108.9 | 1.024 | 0.09 | S19 | 46.7 | 50.5 | 0.248 | 0.037 |
| C20 [c] | 0 | 8.8 | 0.996 | 0.087 | S20 [s] | 4.3 | 19.6 | 0.266 | 0.044 |
| C21 | 766.1 | 746.8 | 1.033 | 0.035 | S21 | 953.9 | 946.3 | 0.24 | 0.01 |
| C22 | 134.2 | 119.6 | 0.982 | 0.055 | S22 | 33.1 | 29.3 | 0.272 | 0.034 |
| C23 | 14.3 | 4.3 | 0.954 | 0.058 | **S23** | **0** | **0** | **0.264** | **0.038** |
| C24 | 101.8 | 96.5 | 1.084 | 0.101 | S24 | 30.1 | 33.9 | 0.302 | 0.057 |
| C25 [c] | 8.3 | 17.1 | 1.028 | 0.094 | S25 [s] | 1.6 | 16.9 | 0.266 | 0.039 |

[c] models failed to converge; [s] singular fit

Appendix S7

**Comparing the cumulative distribution function of subsets of data**

Cumulative distributions of effort-adjusted catch, log($CPUE_C$) [log(individual (trap day)$^{-1}$)], and effort-adjusted species richness, log($CPUE_S$) [log(species (m day)$^{-0.25}$)], for categorical variables (*Unit*, *Season*, *Continent*, *RowWidth*, *Funnel*, *Fluid*) were compared between the groups of records using the Kolmogorov–Smirnov test.

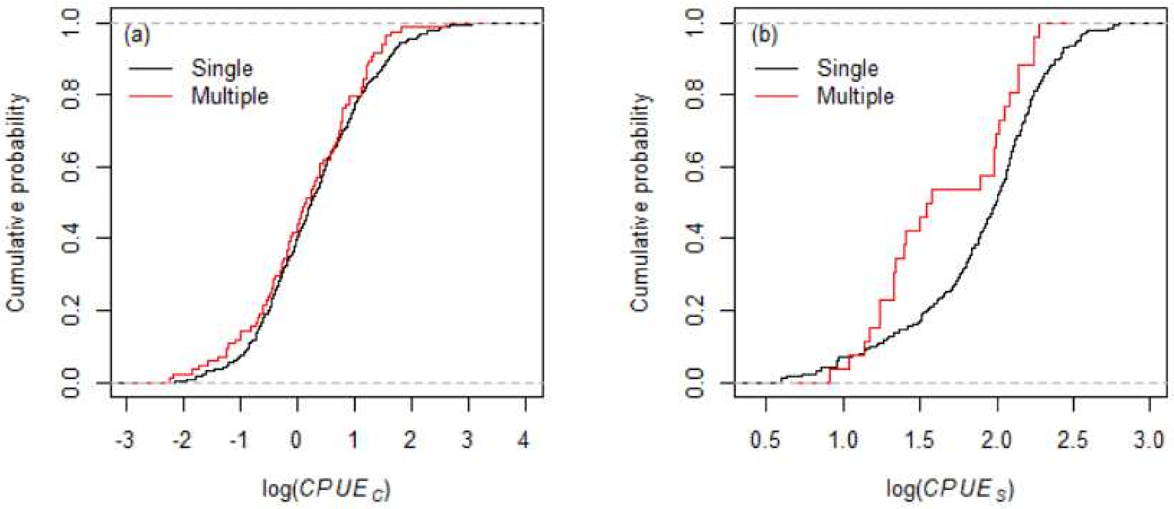


Figure S1 Cumulative distribution functions for subsets of data according to the level of data aggregation over experimental season (single season vs. multiple seasons). (a) log($CPUE_C$): $D$ = 0.077, $P$ = 0.761; (b) log($CPUE_S$): $D$ = 0.328, $P$ = 0.011.

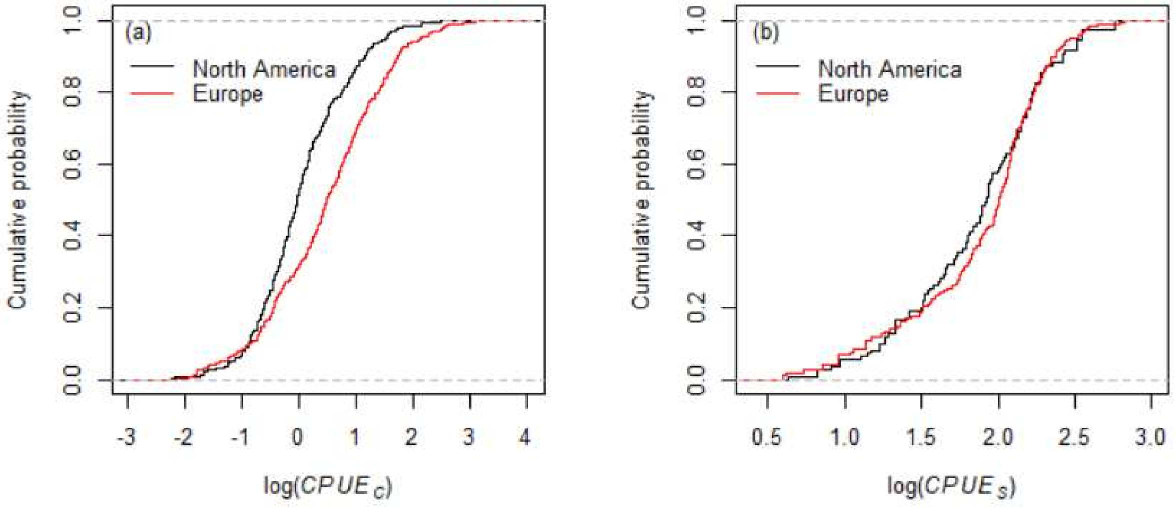


Figure S2 Cumulative distribution functions for subsets of data according to continent of data origin (North America vs. Europe). a) log($CPUE_C$): $D$ = 0.269, $P$ < 0.001; (b) log($CPUE_S$): $D$ = 0.142, $P$ = 0.103.

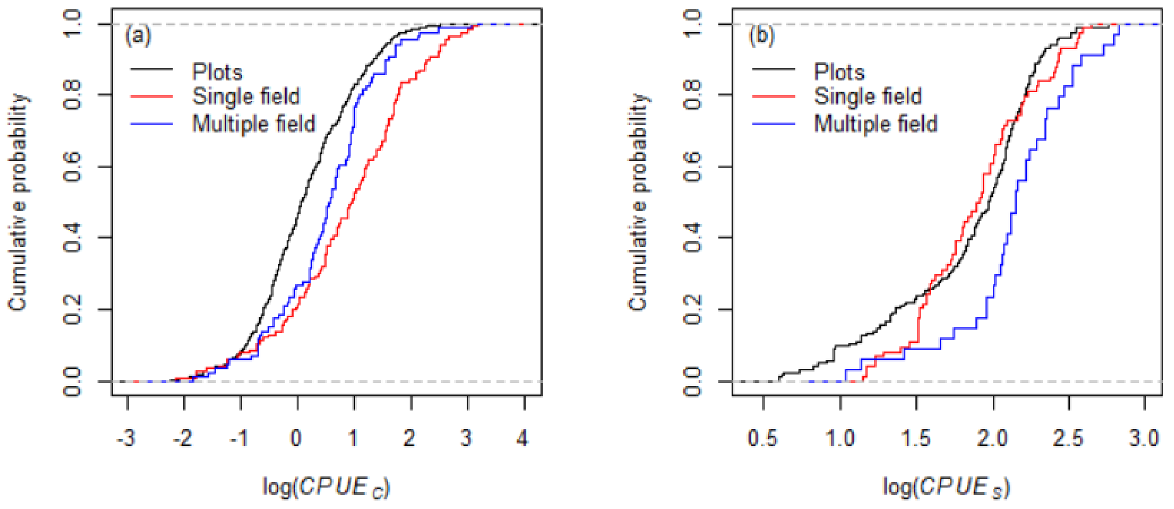


Figure S3 Cumulative distribution functions for subsets of data according to level of data aggregation over experimental units (plots, single field, multiple fields). a) log($CPUE_C$) - plots vs single field: $D$ = 0.347, $P$ < 0.001; plots vs multiple fields: $D$ = 0.282, $P$ < 0.001; single vs. multiple fields: $D$ = 0.259, $P$ = 0.003; (b) log($CPUE_S$) - plots vs single field: $D$ = 0.130, $P$ = 0.304; plots vs multiple fields: $D$ = 0.304, $P$ = 0.009; single vs. multiple fields: $D$ = 0.405, $P$ = 0.001.

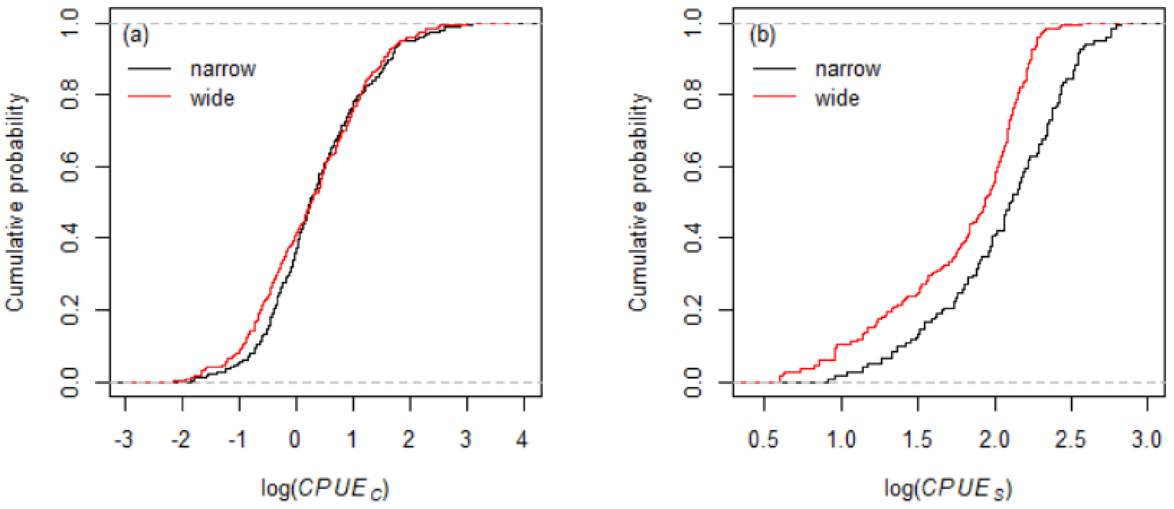


Figure S4 Cumulative distribution functions for subsets of data according to row width (crops with narrow vs. wide rows). a) log($CPUE_C$): $D$ = 0.097, $P$ = 0.067; (b) log($CPUE_S$): $D$ = 0.322, $P$ < 0.001.

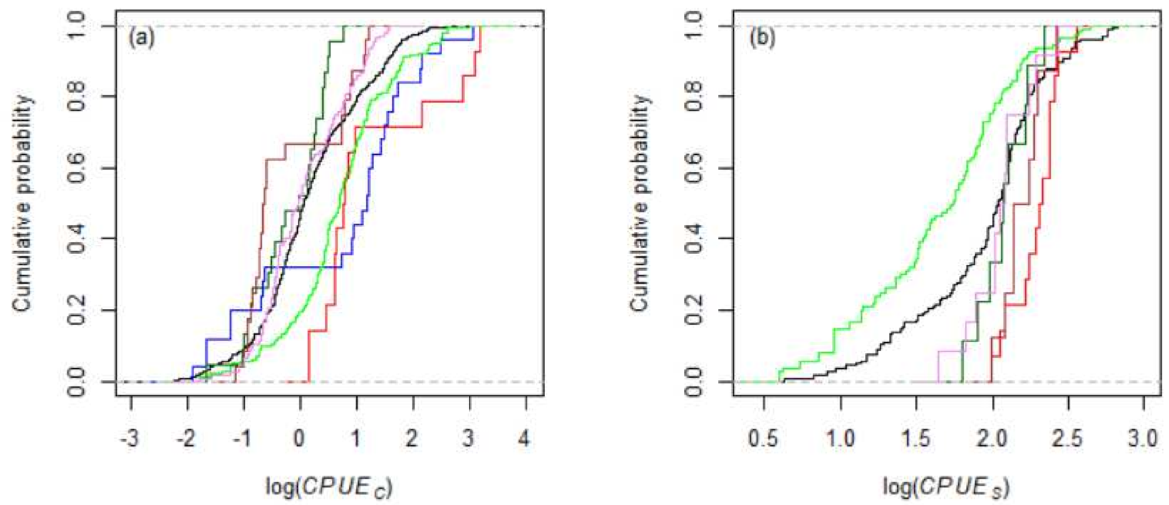


Figure S5 Cumulative distribution functions for subsets of data according to the collecting fluid used in the traps. a) log($CPUE_C$); (b) log($CPUE_S$). Black - antifreeze; red – acetic acid; blue - alcohols; green - formalin; dark green – live traps; violet – water; brown – other fluids. Differences between the distribution functions are shown in Table S1.

Table S1 Differences between the cumulative distribution functions for subsets of data according to the collecting fluid used in the traps compared by a Kolmogorov-Smirnov test. a) log($CPUE_C$); (b) log($CPUE_S$). Above the diagonal – criterion *D*, below the diagonal – *P*-value.

| a) | Antifreeze | Acetic acid | Alcohols | Formaldehyde | Live | Water | Other |
|---|---|---|---|---|---|---|---|
| Antifreeze | - | 0.528 | 0.410 | 0.326 | 0.268 | 0.114 | 0.443 |
| Acetic acid | 0.001 | - | 0.320 | 0.244 | 0.742 | 0.589 | 0.667 |
| Alcohols | <0.001 | 0.317 | - | 0.275 | 0.640 | 0.472 | 0.515 |
| Formaldehyde | <0.001 | 0.433 | 0.079 | - | 0.529 | 0.381 | 0.528 |
| Live | 0.087 | <0.001 | <0.001 | <0.001 | - | 0.246 | 0.364 |
| Water | 0.223 | <0.001 | <0.001 | <0.001 | 0.201 | - | 0.438 |
| Other | <0.001 | <0.001 | 0.003 | <0.001 | 0.089 | 0.001 | - |

b)

| | Antifreeze | Acetic acid | Alcohols | Formaldehyde | Live | Water | Other |
|---|---|---|---|---|---|---|---|
| Antifreeze | - | 0.552 | - | 0.354 | 0.291 | 0.218 | 0.457 |
| Acetic acid | <0.001 | - | - | 0.755 | 0.603 | 0.560 | 0.446 |
| Alcohols | - | - | - | - | - | - | - |
| Formaldehyde | <0.001 | <0.001 | - | - | 0.555 | 0.523 | 0.755 |
| Live | 0.466 | 0.037 | - | 0.012 | - | 0.194 | 0.431 |
| Water | 0.662 | 0.035 | - | 0.005 | 0.990 | - | 0.500 |
| Other | 0.083 | 0.262 | - | <0.001 | 0.412 | 0.181 | - |

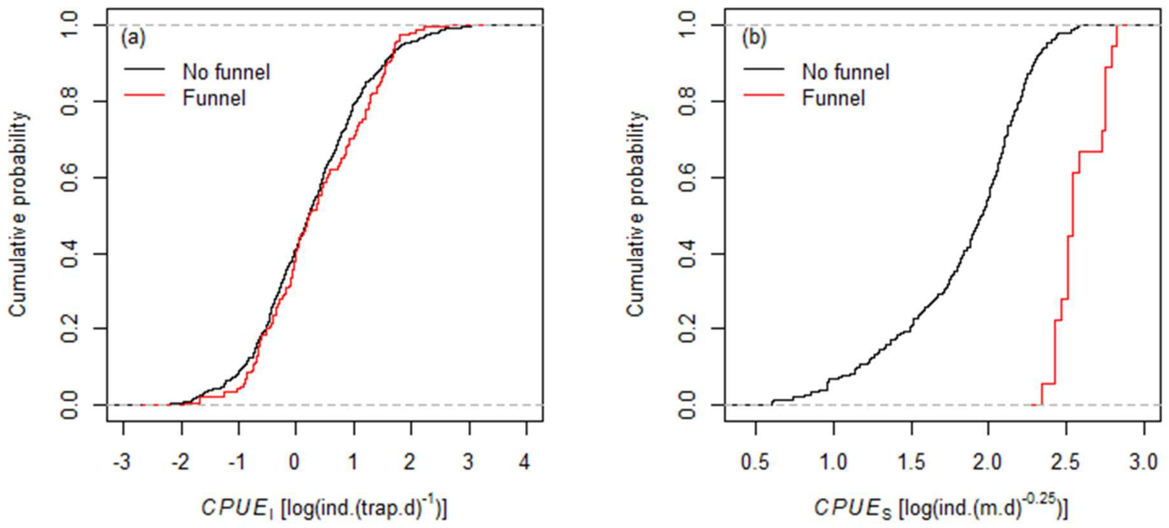


Figure S6 Cumulative distribution functions of *CPUE* for subsets of data according to the use of funnel inside the trap (with or without a funnel). a) $CPUE_C$: $D$ = 0.091, $P$ = 0.302; (b) $CPUE_S$: $D$ = 0.934, $P$ < 0.001.

Appendix S8

**The lists of top models used for model averaging**

Table S1 List of top models from model comparison based on ΔAICc and using dredge function of the R package MuMIn. Only models with ΔAICc < 6 were included. Global models, C36 and S36, for model averaging are defined in Appendix S4, Table S3.

| Fixed effects | df | LL | AICc | ΔAICc | weights | Marginal $R^2$ | Conditional $R^2$ |
|---|---|---|---|---|---|---|---|
| Individuals | | | | | | | |
| *RowWidth + Fluid* | 11 | -621.05 | 1264.47 | 0 | 0.275 | 0.0734 | 0.4451 |
| *RowWidth + Unit* | 7 | -626.17 | 1266.50 | 2.03 | 0.100 | 0.0499 | 0.4414 |
| *RowWidth + Fluid + Funnel* | 12 | -621.13 | 1266.70 | 2.23 | 0.090 | 0.0765 | 0.4483 |
| *RowWidth + Fluid + Unit* | 13 | -620.41 | 1267.33 | 2.86 | 0.066 | 0.0915 | 0.4539 |
| *RowWidth + Fluid + Season* | 12 | -621.47 | 1267.39 | 2.92 | 0.064 | 0.0753 | 0.4462 |
| *Continent + RowWidth + Fluid* | 12 | -621.64 | 1267.73 | 3.26 | 0.054 | 0.0742 | 0.4478 |
| *RowWidth* | 5 | -629.18 | 1268.44 | 3.97 | 0.038 | 0.0162 | 0.4404 |
| *Continent + RowWidth + Unit* | 8 | -626.24 | 1268.68 | 4.21 | 0.033 | 0.0560 | 0.4446 |
| *RowWidth + Funnel + Unit* | 8 | -626.43 | 1269.07 | 4.60 | 0.028 | 0.0523 | 0.4452 |
| *RowWidth + Fluid + Funnel + Unit* | 14 | -620.42 | 1269.44 | 4.97 | 0.023 | 0.0963 | 0.4579 |
| *Continent + RowWidth* | 6 | -628.67 | 1269.46 | 4.99 | 0.023 | 0.0268 | 0.4420 |
| *RowWidth + Fluid + Funnel + Season* | 13 | -621.48 | 1269.47 | 5.00 | 0.023 | 0.0793 | 0.4495 |
| *RowWidth + Season + Unit* | 8 | -626.77 | 1269.73 | 5.26 | 0.020 | 0.0506 | 0.4425 |
| *Continent + RowWidth + Fluid + Funnel* | 13 | -621.70 | 1269.91 | 5.44 | 0.018 | 0.0775 | 0.4512 |
| *RowWidth + Fluid + Season + Unit* | 14 | -620.74 | 1270.08 | 5.61 | 0.017 | 0.0947 | 0.4551 |
| Species richness | | | | | | | |
| *RowWidth* | 5 | -629.18 | 1291.26 | 0 | 0.826 | 0.0162 | 0.4404 |
| *Continent + RowWidth* | 6 | -628.67 | 1296.83 | 5.57 | 0.051 | 0.0268 | 0.4420 |

Table S2 List of top models from model comparison based on ΔBIC and using dredge function of the R package MuMIn. Only models with ΔBICc < 6 were included. Global model for model averaging are listed in Appendix S4, Table S3.

| Fixed effects | df | LL | BIC | ΔBIC | weights | Marginal $R^2$ | Conditional $R^2$ |
|---|---|---|---|---|---|---|---|
| Individuals | | | | | | | |
| *RowWidth + Fluid + Funnel + Season* | 11 | -952.63 | 1928.17 | 0 | 0.226 | 0.2122 | 0.4884 |
| *Continent + RowWidth + Fluid + Funnel + Season* | 12 | -952.34 | 1929.74 | 1.57 | 0.103 | 0.2150 | 0.4875 |
| *RowWidth + Funnel + Season* | 6 | -958.87 | 1930.03 | 1.86 | 0.089 | 0.1516 | 0.5061 |
| *Continent + RowWidth + Funnel + Season* | 7 | -957.85 | 1930.09 | 1.92 | 0.087 | 0.1668 | 0.5056 |
| *RowWidth + Fluid + Funnel + Season + Year* | 12 | -952.63 | 1930.34 | 2.17 | 0.076 | 0.2122 | 0.4884 |
| *RowWidth + Fluid + Funnel + Season + Unit* | 13 | -951.84 | 1930.94 | 2.77 | 0.057 | 0.2189 | 0.4864 |
| *Continent + RowWidth + Fluid + Funnel + Season + Year* | 13 | -952.33 | 1931.90 | 3.73 | 0.035 | 0.2153 | 0.4873 |
| *RowWidth + Funnel + Season + Year* | 7 | -958.87 | 1932.12 | 3.95 | 0.031 | 0.1516 | 0.5061 |
| *Continent + RowWidth + Funnel + Season + Year* | 8 | -957.85 | 1932.19 | 4.02 | 0.030 | 0.1669 | 0.5056 |
| *RowWidth + Funnel* | 5 | -961.14 | 1932.48 | 4.31 | 0.026 | 0.1385 | 0.5266 |
| *Continent + RowWidth + Funnel* | 6 | -960.34 | 1932.96 | 4.79 | 0.021 | 0.1502 | 0.5250 |
| *Continent + RowWidth + Fluid + Funnel + Season* | 14 | -951.78 | 1933.01 | 4.84 | 0.020 | 0.2193 | 0.4859 |
| *RowWidth + Fluid + Funnel + Season + Unit + Year* | 14 | -951.84 | 1933.13 | 4.96 | 0.019 | 0.219 | 0.4866 |
| *RowWidth + Season* | 5 | -961.67 | 1933.54 | 5.37 | 0.015 | 0.0883 | 0.4976 |
| *RowWidth + Funnel + Season + Unit* | 8 | -958.68 | 1933.85 | 5.68 | 0.013 | 0.1531 | 0.5044 |
| *RowWidth + Fluid + Funnel* | 10 | -956.62 | 1933.99 | 5.82 | 0.012 | 0.1991 | 0.5231 |
| *Continent + RowWidth + Funnel + Season + Unit* | 9 | -957.77 | 1934.15 | 5.98 | 0.011 | 0.1677 | 0.5050 |
| Species richness | | | | | | | |
| *RowWidth* | 4 | -963.96 | 1950.80 | 0 | 0.268 | 0.0721 | 0.5211 |
| *RowWidth +Funnel* | 5 | -961.14 | 1950.88 | 0.09 | 0.257 | 0.1385 | 0.5266 |
| *RowWidth + Season* | 5 | -961.67 | 1951.94 | 1.14 | 0.151 | 0.0883 | 0.4976 |

| | | | | | | | |
|---|---|---|---|---|---|---|---|
| *RowWidth + Funnel + Season* | 6 | -958.87 | 1952.07 | 1.27 | 0.142 | 0.1516 | 0.5061 |
| *Continent + RowWidth + Funnel* | 6 | -960.34 | 1955.00 | 4.20 | 0.033 | 0.1502 | 0.5250 |
| *Continent + RowWidth* | 5 | -963.49 | 1955.59 | 4.79 | 0.024 | 0.0854 | 0.5232 |
| *Continent + RowWidth + Funnel + Season* | 7 | -957.85 | 1955.75 | 4.95 | 0.023 | 0.1668 | 0.5056 |
| *RowWidth + Year* | 5 | -963.87 | 1956.34 | 5.55 | 0.017 | 0.0734 | 0.5220 |
| *Continent + RowWidth + Season* | 6 | -961.04 | 1956.40 | 5.60 | 0.016 | 0.1054 | 0.5009 |
| *RowWidth + Funnel + Year* | 6 | -961.13 | 1956.59 | 5.79 | 0.015 | 0.1389 | 0.5264 |
